# Separate and overlapping mechanisms of statistical regularities and salience processing in the occipital cortex and dorsal attention network

**DOI:** 10.1101/2023.09.25.557901

**Authors:** Bertrand Beffara, Fadila Hadj-Bouziane, Suliann Ben Hamed, C. Nico Boehler, Leonardo Chelazzi, Elisa Santandrea, Emiliano Macaluso

**Affiliations:** Université Claude Bernard Lyon 1, CNRS, INSERM, Centre de Recherche en Neurosciences de Lyon (CRNL), U1028 UMR5292, IMPACT, F-69500, Bron, France; Institut des Sciences Cognitives Marc Jeannerod, Lyon, UMR5229, CNRS, Université de Lyon, France; Department of Experimental Psychology, Ghent University, Belgium; Department of Neuroscience, Biomedicine and Movement Sciences, University of Verona, Italy

**Author notes:** <u>Corresponding author:</u> Bertrand Beffara, Lyon Neuroscience Research Center (Impact Team), 16 avenue du Doyen Lépine, 69500 BRON cedex, France.

**Keywords:** Attention, Vision, Space – fMRI, Connectivity

## Abstract

Attention selects behaviorally relevant inputs for in-depth processing. Beside the role of traditional signals related to goal-directed and stimulus-driven control, a debate exists regarding the mechanisms governing the effect of statistical regularities on attentional selection, and how these are integrated with other control signals. Using a visuo-spatial search task under fMRI, we tested the joint effects of statistical regularities and stimulus-driven salience. We found that both types of signals modulated occipital activity in a spatially-specific manner. Salience acted primarily by reducing the attention bias towards the target location when associated with irrelevant distractors, while statistical regularities reduced this attention bias when the target was presented at a low probability location, particularly at the lower levels of the visual hierarchy. In addition, we found that both statistical regularities and salience activated the dorsal frontoparietal network. Additional exploratory analyses of functional connectivity revealed that only statistical regularities modulated the inter-regional coupling between the posterior parietal cortex and the occipital cortex. These results show that statistical regularities and salience signals are both spatially represented at the occipital level, but that their integration into attentional processing priorities relies on dissociable brain mechanisms.

## 1. INTRODUCTION

Visual selective attention (VSA) is a cognitive function that allows us to prioritize behaviorally relevant information among the large amount of visual information that competes for cognitive processing (Chelazzi et al., 2011; Clark, 2003; Dukas, 2004). Traditionally, the factors accounting for the behavioral relevance of visual items - or locations - have been classified in two categories: endogenous and exogenous signals. Endogenous signals refer to information related to the current goals of the participant, while exogenous, stimulus-driven signals, guide selection on the basis of low-level physical features of the sensory input (e.g. color, motion or orientation contrasts, see Desimone & Duncan, 1995; Egeth & Yantis, 1997; Itti & Koch, 2001). This dichotomy has been linked to separate functional networks in the fronto-parietal cortex that are thought to guide attention on the basis of endogenous (dorsal network) or exogenous signals (ventral network; Chica et al., 2011; Corbetta et al., 2000; Corbetta & Shulman, 2002). Nonetheless, this traditional account has been recently challenged with evidence suggesting that endogenous and exogenous signals are not processed by fully independent systems, but rather work together to jointly affect the allocation of attentional resources, leading to different behavioral outcomes depending on the task configuration and on the relative weights of these signals (Anderson, 2021; Beffara et al., 2022; Chelazzi et al., 2019; Luck et al., 2021; Sprague et al., 2018).

In addition, there is now extensive evidence that other types of signals can also contribute to VSA, including emotional content (Todd & Manaligod, 2018; Vuilleumier, 2005), semantics (Gibson & Kingstone, 2006), reward (Anderson, 2016; Anderson et al., 2011; Awh et al., 2012; Bourgeois et al., 2016; Chelazzi et al., 2013, 2014; Pessoa, 2010, 2015) and social factors (Klein et al., 2009). An additional category of signals concerns statistical regularities (Awh et al., 2012). This refers to any recurring pattern of sensory information that can bias attentional selection, regardless of sensory salience and participant’s goals (Awh et al., 2012). Previous studies using visual search tasks revealed that statistical regularities can lead to the facilitation of target selection, but also to reduced processing, depending on the characteristics of the search task (Melloni et al., 2012; Wolfe et al., 2003; see also below). Expectations play a role also in endogenous spatial cueing tasks (Posner, 1980) that systematically show improvements for targets presented at the cued/expected location, as compared with neutral or invalidly cued targets. Nonetheless, the two paradigms (spatial cueing and search) differ in many respects and, in particular, statistical regularities in search tasks imply learning about the probability of spatial distributions over time/trials, while cueing tasks do not rely on any long-lasting changes of spatial representations (i.e. the cued location is typically randomized across trials).

Previous studies using search tasks (e.g. Melloni et al., 2012; Wolfe et al., 2003) highlighted the impact of statistical regularities on behavioral performance, but they did not assess how regularities affect processing at specific spatial locations. Addressing this issue is crucial to understand the relation between statistical regularities and other signals that contribute to assigning spatial processing priorities (e.g. goals and sensory salience; see also the “priority maps framework”, below). With this aim, a behavioral study by Ferrante et al. (2018) tested the joint contribution of target-probability location and distractor’s salience. The results highlighted a significant effect of target-probability location: targets presented at the high target-probability location (HTPL) were associated with better performance than when targets were presented at low target-probability locations (LTPL). In addition, the results showed that the inclusion of a salient distractor in the search display reduced participants’ performance. The interaction between the 2 factors was also significant, with a stronger effect of target-probability location in salient distractor-present compared to salient distractor-absent conditions. These findings demonstrate that statistical regularities and sensory salience can jointly contribute to target selection in the spatial domain.

At the brain level, Melloni et al. (2012) showed that statistical regularities can modulate activity at different levels of the visual hierarchy. The authors compared trials in “fixed” blocks when the search array never contained a salient distractor vs. trials in “mixed” blocks that contained both arrays with and without salient distractors. The imaging results showed that in visual areas V2 and V4 the bias of activity favoring target (vs. distractor) processing was higher in the “fixed” blocks, compared to “mixed” blocks. The whole-brain analyses also revealed activation of the intraparietal sulcus (IPS) in the dorsal fronto-parietal network, related to the interaction between regularities of salient distractor presence and salience, suggesting that IPS is involved in the filtering of salient distractors. In a more recent study, Won et al. (2020) extended this work by testing how regularities of salient distractor’s presence specifically modulate the occipital activity of the salient distractor’s representation. They found that, when the participants were informed about the (high/low) likelihood that the search array would contain a salient distractor, the activity of occipital regions representing the salient distractor was reduced in the “high” compared with the “low” likelihood condition. This suggests that statistical regularities concerning the presence of a salient distractor in the upcoming trial can modulate the activation associated with the distractor processing and, thus, that the high likelihood of salient distractor presence can facilitate distractor suppression. Nonetheless, at odds with the whole-brain results of Melloni et al. (2012), Won et al. (2020) found larger activation of the IPS for salient distractors presented in low vs. high likelihood conditions. Won et al. (2020) suggested that regions of the frontoparietal network were not recruited for distractor suppression (under high distractor singleton likelihood), which instead would occur automatically within the occipital cortex, provided that distractor suppression occurs in an efficient manner (see also Geng & Duarte, 2021, for a review).

In another recent study, Zhang et al. (2021) manipulated the probability for a salient distractor to be presented in a specific spatial location, hence addressing more specifically the role of statistical regularities in the spatial domain. In line with the two previous studies, the results showed that salient distractors presented at the high probability location elicited weaker responses in the corresponding occipital representation, compared with the salient distractors presented at low probability locations. Also, consistent with Won et al. (2020), whole-brain analyses revealed greater activation of the superior parietal lobule (SPL) when the salient distractor was at a low compared to a high probability location. Altogether, these results show that the statistical distribution (presence and/or location) of salient distractors modulates spatially-specific responses in the occipital visual cortex (Melloni et al., 2012; Won et al., 2020; Zhang et al., 2021). This modulatory effect co-occurs with reduced activation of dorsal parietal regions (SPL /IPS) that may mediate distractor suppression when salient distractors are presented at a high probability location (Geng & Duarte, 2021; Won et al., 2020; Zhang et al., 2021).

These findings can be interpreted in the framework of recent models of attention control positing that attentional gains are represented in “spatial priority maps” that would combine different types of control signals (Bisley & Mirpour, 2019; Gottlieb, 2007; Ptak, 2012; Thompson & Bichot, 2005). Priority maps have been associated with regions of the dorsal fronto-parietal attention network (IPS, FEF, Bisley & Mirpour, 2019; Gottlieb, 2007; Ptak, 2012; Thompson & Bichot, 2005), as well as with sub-cortical structures such as the superior colliculus (Bisley & Mirpour, 2019). The priority map framework would predict that statistical regularities should enhance spatial representations of task-relevant target location (e.g. in the occipital cortex) and suppress the location of expected task-irrelevant distractors (see discussion in Zhang et al., 2021, and see behavioral results in Ferrante et al., 2018).

Although this specific prediction has never been formally tested using spatial attention protocols, statistical regularities related to task-relevant targets have been instead associated with so-called “expectation suppression” rather than enhancement (Alink & Blank, 2021; Richter & de Lange, 2019). In particular, Richter & de Lange (2019) used fMRI to investigate the non-spatial responses to predicted/expected stimuli. In a training phase, on each trial a leading image (i.e. the cue) was 100% predictive of the subsequent trailing image (i.e. the target). During the test phase in the scanner, the leading image was now only 50% predictive of the trailing image. The imaging results showed weaker V1 responses when the trailing image followed the predictive leading image than when it followed a non-predictive trailing image (see also Alink et al., 2010). Alink & Blank (2021) proposed that expectations regarding the target could actually be considered as a form of salience modulation, which could explain why expected targets elicited weaker occipital responses: their salience is decreased compared to the unexpected targets.

Accordingly, to date, there is no consensus regarding the mechanisms through which attention integrates target spatial statistical regularities to the spatial priority maps (e.g. see Melloni et al., 2012, who found higher parietal activations for expected display configurations, vs. Won et al., 2020; Zhang et al., 2021, who instead reported decreased parietal activation for expected display configurations). To address this further, there are two main issues that need to be considered. First, do targets presented at a high target-probability location (HTPL) lead to an increase or a decrease of the corresponding representations in the occipital cortex (priority maps vs. expectation suppression frameworks)? And, second, if target-probability location work in a similar manner as sensory salience (see Alink & Blank, 2021), do similar brain mechanisms govern how salience and statistical regularities contribute to instantiating attention spatial priorities?

Here we addressed these issues by directly testing the combined effects of salience and statistical regularities. We employed a visual search task and measured the activation of occipital regions representing the location of the target and of the distractors, as a function of target-probability location and target/distractor salience. In the priority maps framework, we expected increased activation for targets presented in high vs. low probability location (HTPL > LTPL), with sensory salience that should further boost these spatially-specific responses when the target is also the salient item (Awh et al., 2012; Zhang et al., 2021). As an alternative hypothesis, if mechanisms of “expectation suppression” apply also to recurring targets (see Zhang et al., 2021, for the case of recurring distractors), one may predict a reduction of the spatial bias towards the representation of the target location, for targets presented at the high vs. low probability locations (HTPL < LTPL). In addition, on the basis of previous work highlighting the role of parieto-occipital interactions in visuo-spatial attention control (Beffara et al., 2022; Corbetta & Shulman, 2002; Vossel et al., 2012, 2014), additional exploratory analyses tested how statistical regularities and salience affected the inter-regional connectivity between the dorsal parietal cortex and the occipital cortex (see also above, Melloni et al., 2012 vs. Won et al., 2020). In brief, within the priority maps framework (see Bisley & Mirpour, 2019 for a review), we employed a visual search task and manipulated statistical regularities and salience with the aim of testing the hypothesis that these two factors would jointly act within a unified representation of visual space, here targeting quadrant-specific representations in the occipital visual cortex.

## 2. MATERIAL & METHODS

### 2.1. Participants

25 right-handed healthy adults were recruited for the study. They had normal or corrected-to-normal vision, no neurological, psychiatric or cognitive impairments and gave their written informed consent to participate in the study. The study was approved by a national ethics committee in biomedical research (Comité de Protection des Personnes: Sud-Méditerranée II, authorization ID: 2019-A00713-54). 23 participants were included in the final analyses (mean age: 25, range 19-32; 14 females). Two participants were excluded because one participant had excessive head movements (> 3 mm) and one participant failed to maintain central fixation during the main attention task.

### 2.2. Experimental design

The experiment comprised 5 functional imaging runs (4 runs of the main attention task, 10 min each; plus 1 localizer run, 10 min) and one anatomical scan (6 min). Visual stimuli were presented using Cogent Graphics, developed by John Romaya at the Wellcome Department of Imaging Neuroscience, running under MATLAB (The MathWorks, Natick, MA). The stimuli were projected on a screen placed at 90cm from the participant’s eyes (1024 × 768 pixels; projected image size: 31.5cm height × 42cm width).

#### 2.2.1. Localizer task

A localizer task was used to identify the occipital spatial representations of each of the four quadrants of the screen (TL, top-left; TR, top-right; BR, bottom-right or BL, bottom-left), where the stimuli of the main attention task were presented (see below). The localizer stimuli consisted in a moving (28°/s) array of small bars (0.5×0.1° each). The array was only visible in a specific screen location on each stimulation block: a 3×3° square aperture located in one of the four quadrants, with a distance of 7° from the center of the screen. The aperture contained approximately 9 bars at a time. All bars were oriented either horizontally or vertically, but the target stimuli that were right/left tilted (45° from vertical). The target-bars appeared in the aperture at unpredictable times (range 1.08-3.24 sec) and the participants had to report the tilt orientation (right/left) by pressing a response-button with the index/middle finger of the right hand. Participants had to maintain central fixation throughout the localizer scan. The stimuli were presented in each screen quadrant for blocks of 14 seconds, interleaved with 12 seconds of central fixation without any visual stimulation. Each quadrant was stimulated 6 times, in a randomized order.

#### 2.2.2. Main attentional task

The main attentional task was split into 4 fMRI runs, each including 126 trials. Each trial started with a 1000ms warning phase when a central fixation dot (0.5° diameter) was displayed in a light grey color, indicating the imminent presentation of the search array. This was followed by the 4-items display that consisted in 4 oriented bars, one in each quadrant of the screen (eccentricity = 7°, size = 2.0 × 0.5°, see Fig. 1). The search array was displayed for 300ms. Each bar contained a dot either in the top or the bottom part of the bar. The target bar was tilted −25° or 25° from the vertical, while the 3 distractor bars were tilted −25° or 25° from the horizontal. The participants’ goal was to indicate the up/down location of the dot placed inside the target-bar by pressing a response-button with the index/middle finger of the right hand. The 3 distractor bars were fully irrelevant for the task. Participants had up to 3000ms to give a response but were instructed to respond as quickly and accurately as possible. The inter-trial interval was between 3000 and 4000ms (uniform distribution). Participants had to maintain central fixation throughout the fMRI-run.

**Figure 1.**
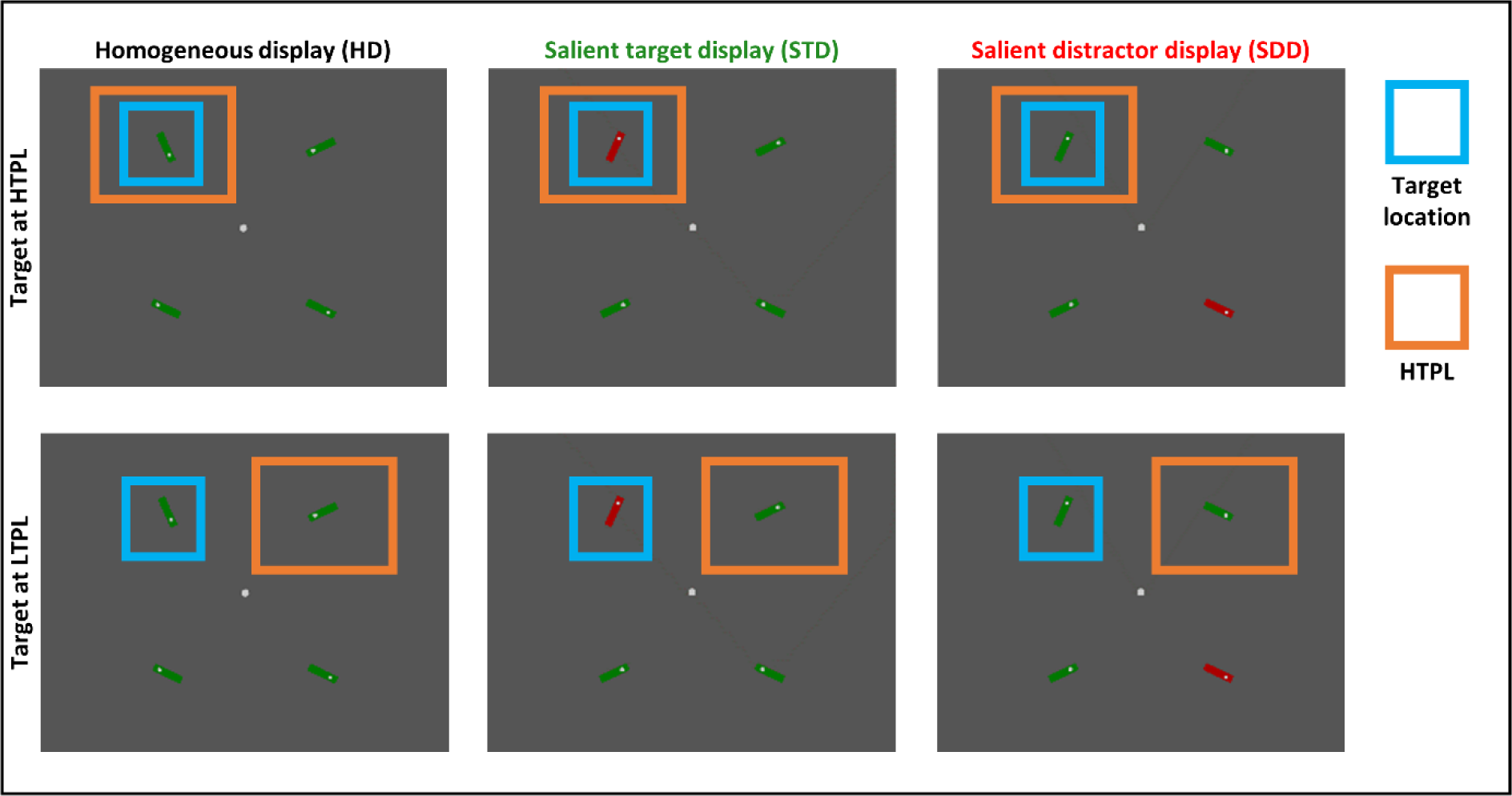
The 6 main experimental conditions. For targets appearing in one specific quadrant (here, top-left), there were 6 possible experimental conditions (hence a total of 4 (quadrants) × 6 (conditions) = 24 conditions). The 6 main experimental conditions correspond to a 2 (statistical regularities of target-probability location conditions) × 3 (salience-conditions) design. Stimulus-driven salience is operationalized using a color singleton item: the 4 stimuli could be of the same color (HD condition), the target could be the color singleton (STD condition) or the distractor diagonal to the target quadrant could be the color singleton (SDD). For each of the 3 salience-conditions, the target (highlighted in blue) could occur either at a high-probability location (HTPL, 50% probability; top panels) or at a low probability location (LTPL, i.e. the 3 remaining quadrants, 16.7% probability for each location, bottom panels).

On each trial, the 4 bars were either all of the same color (green or red, counterbalanced across participants) or included a color singleton (one green bar among three red bars, or vice-versa). This yielded the three salience-conditions that served to operationalize the manipulation of stimulus-driven salience (see Fig. 1): *Homogeneous Display* (**HD**, all stimuli of the same color), *Salient Target Display* (**STD**, when the target was the color singleton) and *Salient Distractor Display* (**SDD**, when a distractor stimulus was the color singleton). In order to reduce the total number of possible conditions, in the SDD condition the salient distractor always appeared in the quadrant located diagonally to the target quadrant. The three salience-conditions were presented in randomized order and with equal probability.

To address the effect of statistical regularities, we manipulated the probability of the target to be presented in a given screen quadrant. In each fMRI-run, the target stimulus was presented with high probability in one of the 4 quadrants (**HTPL**, 50%; Fig. 1, top panels) and with lower probability in the other 3 quadrants location (**LTPL**, 16.67% for each quadrant; Fig. 1, bottom panels). The HTPL quadrant was pseudorandomly assigned to a different quadrant across the 4 fMRI-runs. The combination of salience-condition, target-probability location, and target-location resulted in a fully-factorial design: 3 salience-conditions (HD, STD, SDD) × 2 target-probability location (HTPL, LTPL) × 4 target locations (i.e. the screen quadrants: TL, TR, BR, BL).

### 2.3. Eye tracking

Eye-position was monitored throughout both the localizer scan and the 4 fMRI-runs of the main experiment. The participants’ gaze was tracked using a MR-compatible EyeLink 1000 (SR Research Ltd., Mississauga, Ontario, Canada) at a sampling rate of 500Hz. Calibration was performed using the four quadrants (7° eccentricity), plus the screen center. Eye-tracking data analysis was performed using custom MATLAB scripts. Data were extracted in a 2500ms window starting 500ms before the warning cue. Data were baseline-adjusted using the median of the vertical and horizontal eye-position during the first 500ms. We assessed the quality of each trial’s data considering the percentage of data-points larger than 10°/smaller than −10°, caused by eye blinks or noisy signal. Only trials with less than 50% of these unreliable data-points underwent further analyses (86% of the trials). In these trials, any displacement of eye-position larger than 0.5° and lasting for at least 100ms was classified as a change of fixation. Trials including any fixation outside a central box of 2° from the center of the screen were considered trials with a loss of central fixation (10.4% of the trials with reliable data). These trials were excluded from behavioral analyses and were modeled in a dedicated regressor of non-interest in the imaging analyses.

### 2.4. Behavioral analyses

At the end of the experiment, participants were asked whether they had noticed anything particular during the experiment. They were then asked whether they had noticed any pattern or regularity of the target location during the fourth fMRI-run, and then asked to guess what was the location where most of the target were presented in this last run. Finally, they were asked to guess the sequence of high target-probability locations across the 4 runs.

We analyzed the response times (RT) and accuracy data for the main attentional task using linear mixed models implemented in R studio (Bates et al., 2015). For the RT analysis, trials with wrong/no/late responses and/or including losses of fixation (cf. above) were discarded from the analysis. The model included the log-transformed RTs as the dependent variable and the salience-condition (with 3 levels: HD, STD & SDD), the target probability location condition (with 2 levels: HTPL, LTPL) and their interaction as the explanatory variables. For the accuracy analysis, only trials including losses of fixation were discarded. The model included the response accuracy (correct or wrong) modeled using a binomial law as the dependent variable, and the salience-condition, target-probability location and their interaction as explanatory variables. Because of the repeated measures, both models also included subject-specific intercepts.

### 2.5. Image acquisition and preprocessing

T2*-weighted echoplanar images (EPI) with blood oxygen level-dependent (BOLD) contrast (interleaved multiband sequence, Multiband Factor = 2, 50 slices covering the entire brain, field of view = 220×210.4mm, Repetition Time = 1.72s, echo time = 30ms, phase encoding direction = antero-posterior, slice orientation = approx. axial, voxel size = 2.4 × 2.4 × 2.4mm^3) were obtained using a 3T MRI System (Trio, Siemens). Note that the EPI sequence enabled us to reduce the matrix size in the phase encoding direction (here 95.7%), hence the particular dimensions of the field of view. A high-resolution anatomical scan was acquired using a standard T1-weighted 3D MP-RAGE sequence (Repetition Time = 3s, echo time = 3.8ms, voxel size = 1 × 1 × 1mm^3).

The functional data were preprocessed and analyzed with Statistical Parametric Mapping software SPM12 (Wellcome Department of Imaging Neuroscience, University College London, UK; http://www.fil.ion.ucl.ac.uk/spm). After discarding the first four volumes of each run, images were corrected for head movements. Slice-acquisition delays were corrected using the middle slice as reference. All images were normalized to the SPM12 Tissue Probability Map and re-sampled to 2mm isotropic voxel size. Unsmoothed data were used for the ROI analyses and smoothed data were used for the whole-brain and psychophysiological interactions analyses.

### 2.6. Single-subject analyses

#### 2.6.1. Localizer and individual Regions Of Interest (ROIs)

The procedure for the ROIs definition has been conducted as specified in detail in Beffara et al. (2022). In brief, the single-subject models (GLM) included the 4 conditions corresponding to the 4 stimulated quadrants (i.e. TL, TR, BR, BL locations, blocks of 14 seconds), plus the 6 movements parameters as regressors of non-interest. The regressors of interest were convolved using the SPM12 “Hemodynamic Response Function”. For each participant, we tested the main effects of the side of the visual stimulation at the whole-brain level (e.g.: “TL+BL > TR+BR”, for left hemifield quadrants) and separated voxels responding to stimulation of the upper or the lower quadrant by using inclusive masking with the relevant effect of top/bottom stimulation (e.g. top-left quadrant: “TL > BL”, and bottom-left quadrant: “BL > TL”). Contrasts were thresholded at p-uncorrected = 0.005.

These subject- and quadrant-specific responses were then split based on the Automated Anatomical Labeling atlas (AAL, Tzourio-Mazoyer et al., 2002) in order to define ROIs on the basis of both quadrant-specificity and anatomically-defined occipital Brodmann areas BA17, BA18 and BA19. This resulted in a total of 12 ROIs per participant (i.e. 3 BA: BA17, BA18, BA19 × 4 quadrants: TL, TR, BR, BL). The spatial localization of the 12 quadrant-specific ROIs is shown in Fig. 2. The average size of the ROIs was (mean number of voxels ± SEM): 83.68 ± 4.64 for BA17, 220.58 ± 8.47 for BA18 and 215.23 ± 10.29 for BA19. The subdivision of the quadrant-specific responses in ROIs belonging to BA17/18/19 provided us with a way to operationally define different levels of the visual processing hierarchy (lower-to-higher). Nonetheless, it should be stressed that the current approach did not allow us to specifically identify voxels belonging to the different retinotopic areas (V1, V2, V3, etc.), which constitute the functional hierarchy of the visual system and that do not strictly correspond to the Brodmann areas (see also Fig. S2). Instead, our localizer task was only used to identify, in each participant, the voxels responding specifically to the visual locations where the stimuli were then presented during the main attention task (see below).

**Figure 2.**
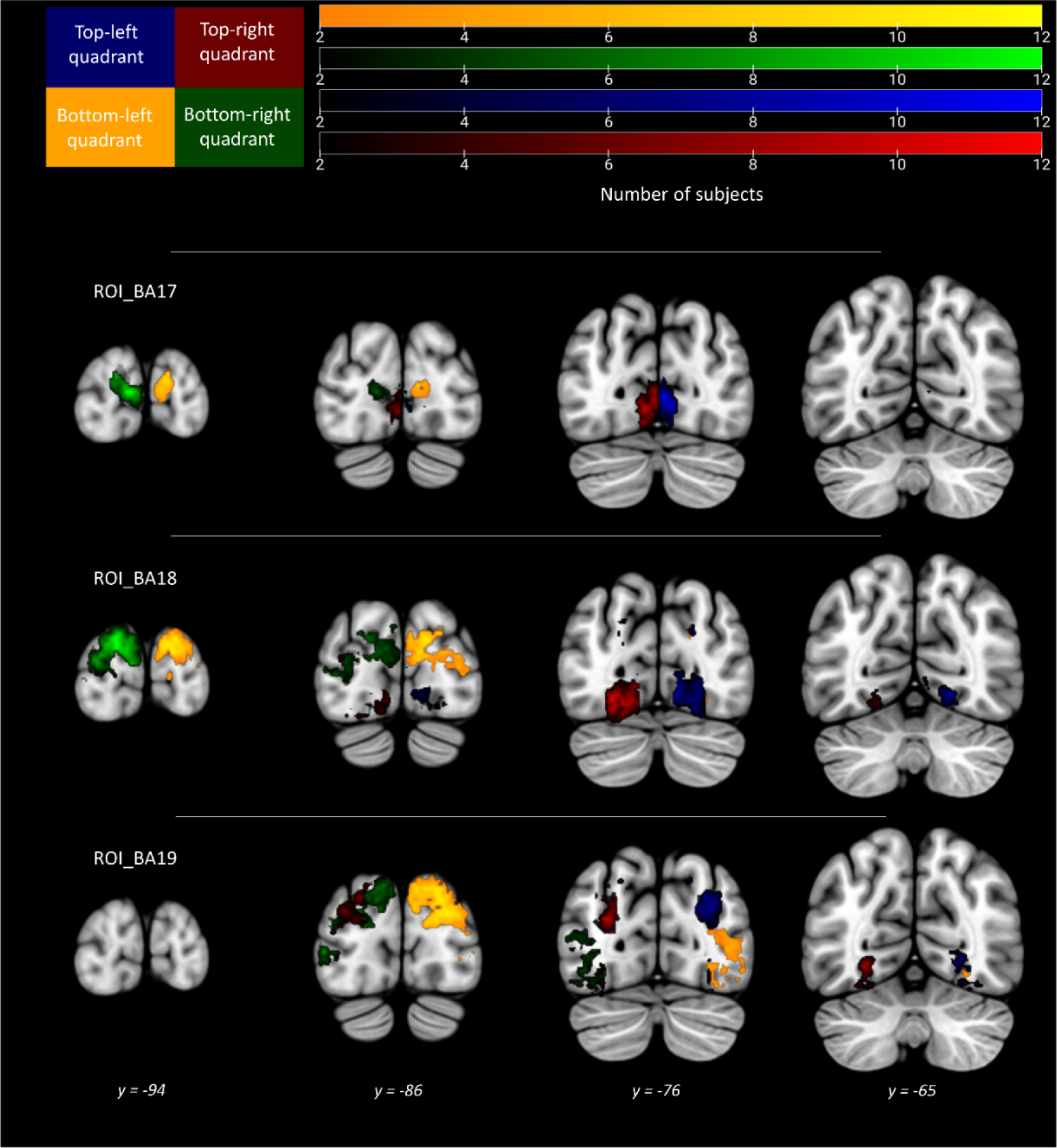
Subject- and quadrant-specific regions of interest in ROI_BA17/18/19. Each color corresponds to voxels responding to a given quadrant of the visual display (cf. top-left schema). The color intensity of each voxel represents the number of subjects responding to the localizer stimuli.

#### 2.6.2. Main task

The main attentional task was analyzed using two separate single-subject GLMs: the first one aimed at investigating quadrant-specific responses in the occipital visual cortex (cf. subject-specific ROI analyses), while the second aimed at testing at the whole-brain level the effects of salience and target-probability location, irrespective of target-quadrant.

In order to study the quadrant-specific responses, the first GLM comprised 24 conditions of interest given by the 2 × 3 × 4 factorial design: target-location probability (HTPL, LTPL) × salience-conditions (HD, STD, SDD) × target-quadrant (TL, TR, BR, BL), plus 8 predictors of non-interest: one regressor including all trials when participants made response errors (no response, incorrect response or response time outside of the 200-3000 ms response window), one regressor including fixation-loss trials and the 6 movement parameters. Each trial was modeled using the canonical Hemodynamic Response Function in SPM12, the event-onsets were time locked to the presentation of the search array and the event-duration was 300ms. Only the parameter estimates of the 24 conditions of interest were used for the subsequent group-level analyses.

The second single-subject GLM aimed at assessing the effects of statistical regularities and salience at the whole-brain level, and comprised 6 conditions of interest: 2 target-probability location (HTPL, LTPL) × 3 salience-conditions (HD, STD, SDD), plus the 8 regressors of non-interest detailed in the first GLM description (see previous paragraph). Each trial was modeled using the canonical Hemodynamic Response Function in SPM12. The event-onsets were time locked to the presentation of the search array and the event-duration was 300ms. Only the parameter estimates of the 6 conditions of interest were used for the subsequent group-level analyses.

### 2.7. Group-level analyses of spatially-specific occipital responses

#### 2.7.1. Target selection, salience and target-probability location

We were interested in the modulation of spatially-specific occipital responses by statistical regularity and visual salience (cf. Fig. 2). For this, we first examined the activity in the ROIs representing the target-quadrant and the diagonal quadrant (see Beffara et al., 2022), as a function of the priority signals present in these two quadrants. To do so, we averaged the response estimates in the occipital ROIs (separately for BA17, BA18 and BA19), considering whether the represented quadrant included the target (“ROI_IN”) or the location diagonal to the target (“ROI_DIAG”, where the salient distractor was presented in the SDD condition). This was done separately for the 3 salience-conditions (HD, STD, SDD) and according to whether the target-quadrant was the high-probability target-location (HTPL) or not (LTPL). This resulted in a single value per subject for each ROI-type (IN/DIAG), BA and experimental condition (3 salience-conditions × 2 statistical regularities conditions). A 2×2×3×3 repeated-measure ANOVA with the factors: quadrant-type (ROI_IN, ROI_DIAG) × target-probability location (HTPL, LTPL) × salience-condition (HD, STD, SDD) × BA-area (BA17, BA18, B19) was used to assess the effects of statistical regularities and visual salience, and their interaction, on the quadrant-specific responses of the 3 BA-areas.

#### 2.7.2. Spatial bias vectors

In addition to considering activity only in the occipital regions representing the target quadrant (ROI_IN) and the opposite (diagonal) quadrant (ROI_DIAG) we examined also how target selection, salience and target-probability location jointly affected the overall 2D representation of visual space, now considering also the activity in the ROIs representing the quadrant ipsilateral to the target quadrant (“ROI_IPSI”, same hemifield as the target, but different up/down location) and the quadrant contralateral to the target (“ROI_CONTRA”, opposite hemifield, but same up/down location as the target). In the priority maps framework it should be considered that all the stimuli present in the visual field concurrently contribute to the distribution of processing properties (Awh et al., 2012; Bisley & Mirpour, 2019; Itti & Koch, 2001) and that stimuli at one location can lead to spatial biases beyond the very specific location of the stimulus (e.g. nearby locations or locations in the same visual hemifield, Gaspelin et al., 2017). In a previous study (Beffara et al. 2022), we proposed a method to investigate the BOLD correlates of these distributed effects of attentional signals across the visual field. These “bias vector analyses” combine the BOLD activity measured in the ROIs that represents the four visual quadrants using vectors’ summation (see Fig. S1). This provides us with a quantitative measure of the direction and strength of the spatial bias in the 2D space, which can then be compared across experimental conditions. In brief, for each of the 24 conditions (see above) and separately for each subject and BA, the mean beta-values of the 4 relevant ROIs (TL/TR/BL/BR) are projected in a 2D plane. Specifically, the mean beta-value of each ROI will determine the length of one vector extending along one of the four 45° diagonals of the 2D-plane (see central panels of Fig. S1). These four vectors are then summed to obtain a condition-specific vector, for which amplitude and direction represent the spatial bias that the relevant condition induces in the BA/visual representation (gray vector in the bottom-right panel of Fig. S1). To compute the bias vectors associated with the 6 conditions of interest (3 salience-conditions × 2 target-probability location conditions), the conditions-specific vectors associated with the four target locations are projected to a single quadrant of the 2D plane (vectors displayed in dotted lines in the lower-left quadrant of Fig. S1, bottom-left panel) and are summed to obtain the “spatial bias vector” associated with the condition of interest (black vector, in the bottom-left panel).

To investigate the effect of the 6 main experimental conditions, for each participant, the data analyses considered the Euclidian distance between the spatial bias vectors and a point positioned on the diagonal corresponding to the location of the target in the frame of reference of the bias vectors (ROI_IN, 45° in the top-left quadrant, see also dotted lines in Fig. 5, top panels). The point was located at the coordinates (−11; 11) that correspond to the maximum absolute value of all X and Y coordinates, across all subjects, conditions and BAs. Any increase of distance between this point and the subject-specific vector expresses a modification of the spatial bias away from the target location (i.e. a decrease of the vector’s magnitude and/or a deviation of the vector’s sense away from the target location, see Fig. 5). The effect of target-probability location and salience-condition on these distance-values were tested using repeated-measures ANOVAs and paired t-tests. Again, it is important to emphasize that any change of the vector bias (e.g. a decrease associated with a specific condition) reflects a modification of the overall representation of space, which can be driven by changes of activity in any (or all) of the 4 visual regions that are used to compute the vectors. For example, a reduction of the bias towards the target location could be driven by less activity in the ROI_IN (i.e. the region that represents the target location), more activity in the ROI_DIAG (i.e. the region representing the location diagonally opposite to the target), but also by differential activity in the ROIs that represent the quadrants ipsilateral and contralateral to the target location (ROI_IPSI and ROI_CONTRA). The latter would shift the vector away from the 45° diagonal and from the (−11, 11) reference point, thus determining a distributed pattern of activity that is less biased towards the actual target location representation.

### 2.8. Whole-brain activations

Beside the modulation of activity in the occipital regions that represent the different visual quadrants (cf. section above), we also examined the effect of target-probability location and salience-condition at the whole-brain level, now irrespective of target location (see also Won et al., 2020; Zhang et al., 2021). The 6 contrast images resulting from the corresponding single-subject GLMs (2 target-probability locations × 3 salience-conditions, see above) were entered in a group-level repeated-measures ANOVA. We then performed the relevant T-contrasts to highlight activations corresponding to the two different types of priority signals (statistical regularities: HTPL > LTPL and LTPL > HTPL; and salience: STD > SDD and SDD > STD). Significant activations reported here were thresholded at p-FWE-corrected = 0.05 at the cluster level, with the cluster-sizes defined at a voxel-wise threshold of p-uncorrected = 0.005, considering the whole brain as the volume of interest. Results are visualized using SPM and MRIcron.

### 2.9. Psychophysiological interactions (PPI)

Our whole-brain analysis (cf. above) revealed overlapping effects of both statistical regularities and salience in the posterior parietal cortex (PPC; see Fig. 6). We sought to further investigate the possible role of the PPC using exploratory analyses of inter-regional connectivity (Friston et al., 1997). The gPPI toolbox (“generalized psycho-physiological interaction”, McLaren et al., 2012) was used to test for condition-dependent changes of connectivity with the PPC. The seed region comprised the voxels showing an effect of both target-probability location (LTPL > HTPL) and salience-conditions (SDD > STD), and that were located within the anatomically AAL-defined BA7 (Tzourio-Mazoyer et al., 2002). The final PPC seed region included 187 voxels. It should be noticed that the overlap between statistical regularities and salience in PPC does not imply that the same population of neurons coded for the two types of signals. Nonetheless, the choice of using these voxels for the analysis of connectivity ensured that the seed ROI included voxels with a consistent pattern of response, related to both salience and statistical regularity. The first-level PPI model included the 6 experimental conditions (2 target-probability locations × 3 salience-conditions), the subject-specific time-series of the PPC seed region, the 6 PPI regressors corresponding to the interactions between each experimental condition and the seed region, plus the 6 movement parameters.

The group analysis comprised a repeated-measures ANOVA with the 6 relevant PPI interaction-terms. We tested for the changes of the PPC connectivity as a function of statistical regularities (LTPL > HTPL) and salience (SDD > STD). The PPI results were first assessed considering the whole-brain as the search volume (p-FWE-corrected = 0.05 at the cluster level, with the cluster-sizes defined at a voxel-wise threshold of p-uncorrected = 0.005). Because of our specific interest in the possible role of PPC in contributing to the effects that we observed in the occipital cortex, we also carried out additional analyses that considered the anatomically-defined occipital ROIs (BA17, BA18, BA19) as volumes of interest, using the Small Volume Correction procedure in SPM12 (Worsley et al., 1996).

## 3. RESULTS

### 3.1. Behavioral data

None of the participants reported having noticed the manipulation of the target-probability location when asked if they noticed something peculiar during the experiment. When asked about the target location imbalance during the last fMRI-run, 7 participants declared that they noticed the imbalance. When asked to guess the HTPL in the last fMRI run, 14 participants were able to correctly report the HTPL. 12 participants declared that they noticed changes of HTPL across runs. Only 2 participants out of 23 were able to report the correct sequence of HTPL across the 4 runs. Only one participant both declared that he/she noticed a change of HTPL across runs and reported the correct HTPL sequence across runs. These data suggest that participants registered the manipulation of target-probability location, but that it is unlikely that they made use of this information to explicitly/voluntarily control attention during the task.

The analysis of the RTs revealed a significant main effect of statistical regularities, with slower RTs in LTPL than in HTPL (F(1,9852) = 98.36, p < 0.001; HTPL, 728 ± 3.09ms; LTPL, 759 ± 3.09ms; fixed-effect Cohen’s d for the HTPL vs. LTPL comparison = 0.16) and a significant main effect of salience, with slower RTs in SDD than in HD, and in HD than in STD (F(2,9852) = 203.36, p < 0.001; HD, 744 ± 3.77ms; STD, 702 ± 3.43ms; SDD, 786 ± 4.02ms; fixed-effect Cohen’s d for the STD vs. SDD comparison = 0.43), see Fig. 3A. The RTs analysis did not reveal any significant interaction between the two factors (F(2,9852) = 0.22, p = 0.80). Accuracy analyses showed a significant main effect of statistical regularities, with lower accuracy in LTPL than in HTPL (χ²(1,10549) = 5.93, p = 0.01; accuracy for: HTPL, 94 ± 0.6%; LTPL, 93 ± 0.7%; odds-ratio for HTPL vs. LTPL = 1.21) as well as a significant main effect of salience, with lower accuracy for SDD than HD and for HD than STD (χ²(2,10549) = 13.22, p = 0.001; accuracy for: HD, 93 ± 0.8%; STD, 94 ± 0.8%; SDD, 92 ± 0.8%; fixed-effect odds-ratio for STD vs. SDD = 1.42), see Fig. 3B. Here also, no significant interaction was found between statistical regularities and visual salience (χ²(2,10549) = 0.02, p = 0.99).

**Figure 3.**
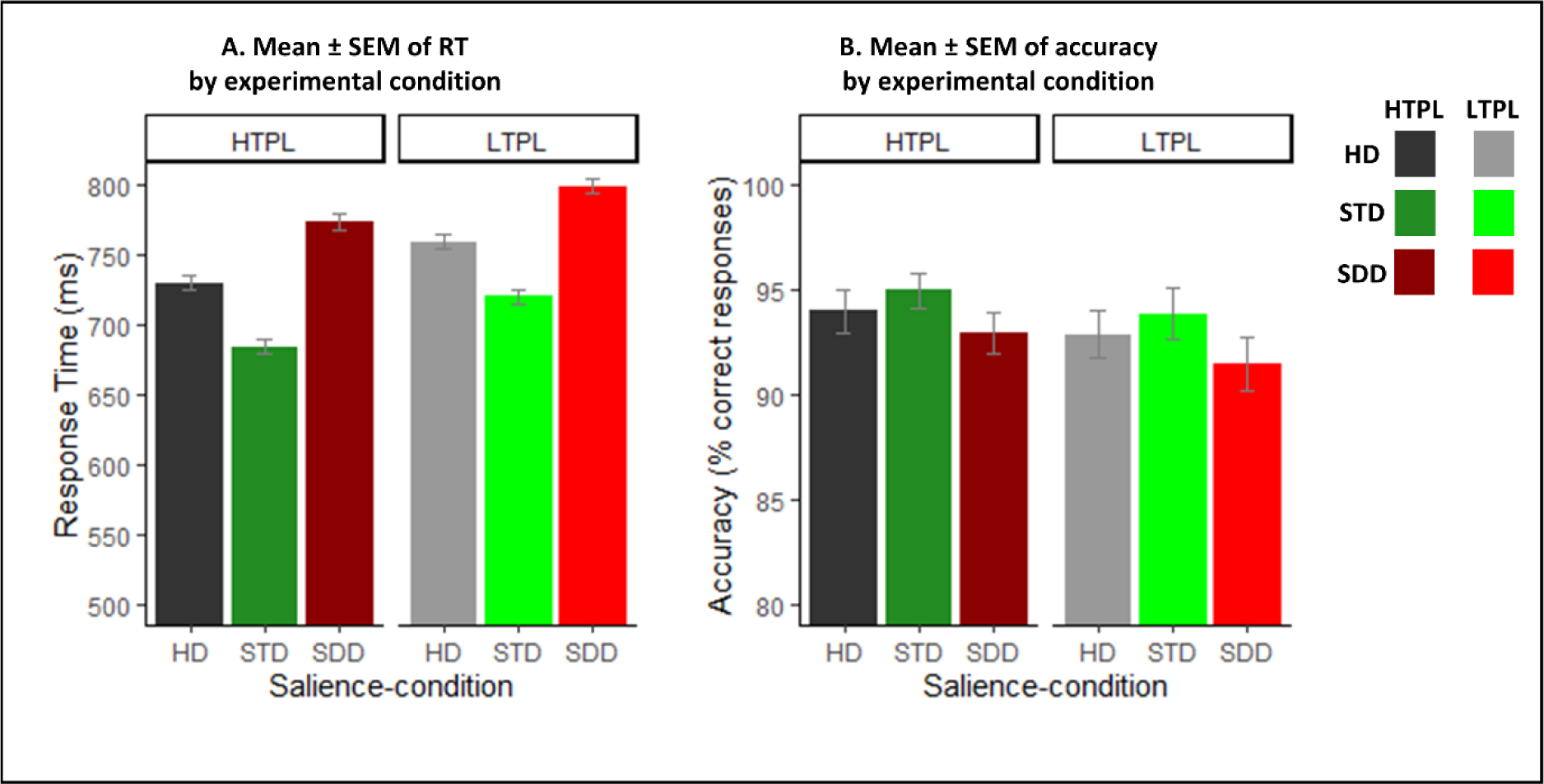
Behavioral data for the 6 main experimental conditions. **A.** The RTs (mean ± SEM) showed significant effects of both salience-condition (STD < HD < SDD) and of target-probability location (HTPL < LTPL), without any interaction between the two factors. **B.** The accuracy data (mean ± SEM) revealed an analogous pattern of results, with significant effects of salience-condition and target-probability location, but no interaction.

The behavioral data demonstrate that target salience facilitates target selection, while distractor salience reduces search performance; and that target-probability location also affects performance, facilitating target selection at the high-probability location compared with performance at the low-probability locations.

### 3.2. Attention priority signals modulate activity in the occipital cortex

#### 3.2.1. Regions representing the target location (ROI_IN) and the diagonal quadrant (ROI_DIAG)

Our first aim was to assess how multiple priority signals present in the visual field modulate the response of quadrant-specific spatial representations in the occipital cortex (Fig. 4). First, we considered activity of the occipital regions representing the current target location (ROI_IN) and the diagonal quadrant, where the salient distractor was presented in the SDD condition (ROI_DIAG). The corresponding 2×3×2×3 ANOVA (ROI_IN/ROI_DIAG × salience-condition × target probability location × BA) revealed significant main effects of ROI-type (F(1,818) = 99.73, p < 0.001, η²p = 0.11), salience-condition (F(2,818) = 6.14, p < 0.002, η²p = 0.02) and BA (F(2,818) = 454.21, p < 0.001, η²p = 0.54). The ROI-type interacted both with BA area (F(2,818), p < 0.005, η²p = 0.01), corresponding to increasingly larger effects of target selection along the visual hierarchy (“ROI_IN > ROI_DIAG”: in BA17, t(137) = 5.46, p < 0.001, Cohen’s d = 0.46; BA18, t(137) = 13.48, p < 0.001, Cohen’s d = 1.15; and BA19, t(137) = 15.79, p < 0.001, Cohen’s d = 1.34), and with the salience-condition (F(2,818), p < 0.01, η²p = 0.0007). The latter corresponds to the larger effect of presenting salient distractors in the ROI_DIAG (“SDD > STD”, t(137) = 6.69, p < 0.001, Cohen’s d = 0.57), compared with salient targets in the ROI_IN (“STD > SDD”, p > 0.9). In fact, in ROI_IN the level of activity was often - and unexpectedly - numerically larger in HD than STD, although this difference did not reach statistical significance in any of the BAs (p > 0.5 in BA17, p > 0.1 in BA18, p > 0.15 in BA19; see also Discussion section concerning the lack of salience effects in the occipital regions representing the task-relevant targets).

**Figure 4.**
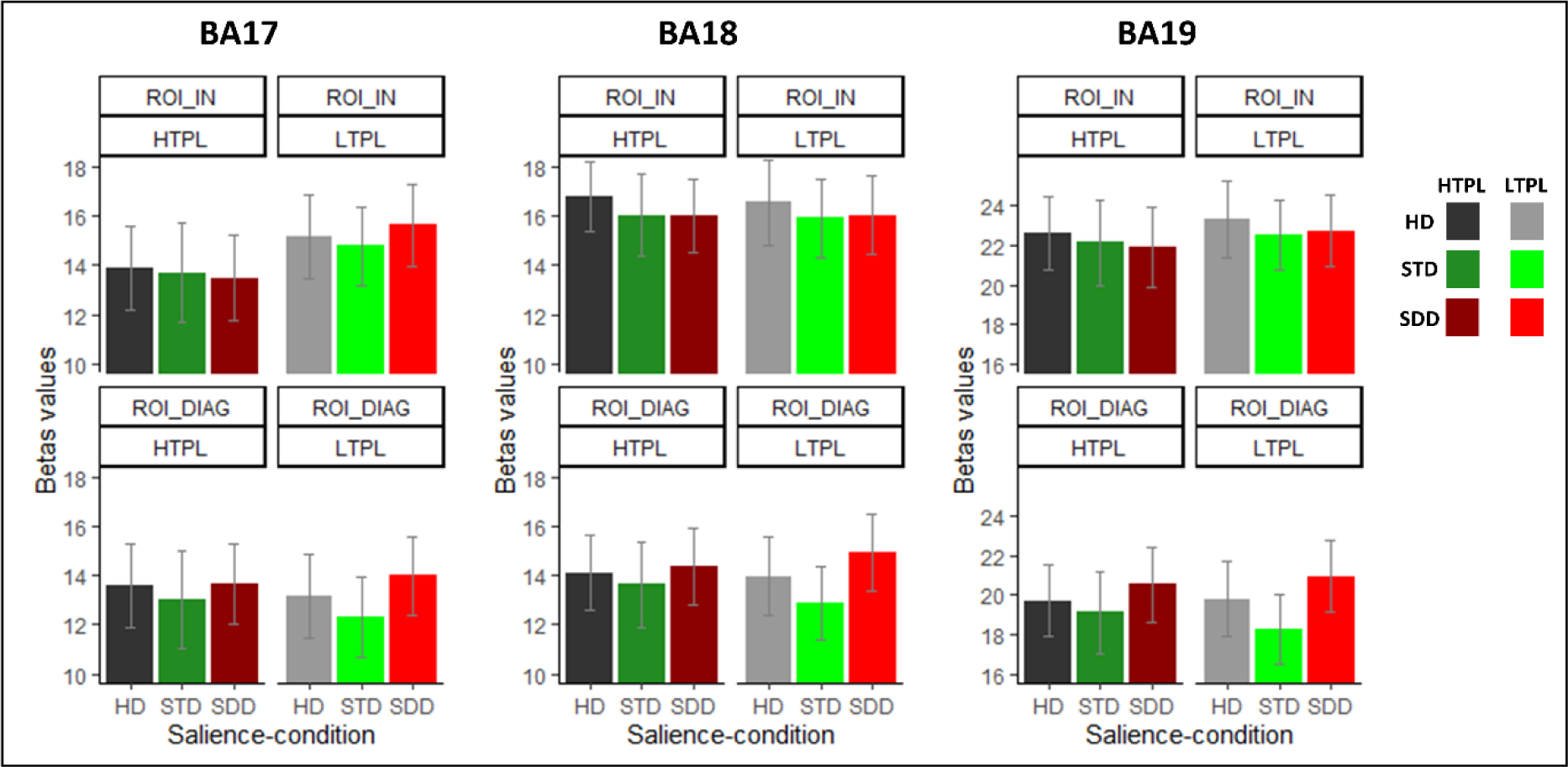
Local responses in ROI_IN and ROI_DIAG as a function of salience-conditions and target-probability location. Left panel shows responses in BA17, middle panel shows responses in BA18 and right panel shows responses in BA19. Grey colors are used for responses in HD, green colors for STD and red colors for SDD. Darker colors are for HTPL, brighter colors are for LTPL. The beta values are expressed in arbitrary units, and their magnitude depends on the scaling of the single-subject design matrix (general linear model) and the data pooling across the target-quadrant (TL, TR, BR, BL; see also Methods section).

The ANOVA also revealed an interaction between ROI-type and target-probability location. Specifically, the ROI_IN activity was larger in LTPL compared with HTPL (“LTPL > HTPL”, t(206) = 3.04, p < 0.005, Cohen’s d = 0.21), while target-probability did not affect activity in ROI_DIAG (“LTPL > HTPL”, p > 0.9). All other effects were non-significant (p > 0.2).

The effect of target-probability location did not interact with the factor BA (p > 0.2), suggesting analogous effects across the 3 areas. Nonetheless, for completeness (see also results of the bias vector analysis below), we tested the effect “LTPL > HTPL” in ROI_IN separately for the 3 BAs. This revealed a significant effect of target-probability location in BA17 (“LTPL > HTPL”, t(68) = 3.42, p < 0.003, Cohen’s d = 0.41), while this effect did not reach significance in BA18 (p > 0.9) and BA19 (p > 0.2).

In sum, we found a significant effect of target selection, with larger activation of the target representation (ROI_IN) than the diagonal distractor representation (ROI_DIAG). This effect was progressively larger from BA17 to BA19. The main finding of the analysis was that salience and statistical learning both affected activity in the occipital cortex, and did so in a spatially-specific manner. We found that salience was most effective when associated with a task-irrelevant distractor (compared with relevant targets) and, conversely, target-probability location impacted most on the target representation, with smaller activation for targets presented at the high compared to the low probability location, especially in BA17. These results demonstrate that both salience and statistical regularities modulate the representation of space in the occipital visual cortex.

#### 3.2.2 Modulation of 2D occipital spatial representations: spatial bias vector analysis

The analyses concerning the local activity in areas representing the target and the diagonal distractor (ROI_IN and ROI_DIAG) revealed effects of salience and statistical regularities, but it should be noted that these analyses did not consider the visual representation of the whole display. To do this, we considered a complementary analysis approach that takes into account the activity of all 4 ROIs that represent the 4 quadrants containing the search items (cf. methods section, and Beffara et al., 2022). Using a 3×2×3 ANOVA we examined the effects of salience-condition, target probability location and BA on the spatial bias vectors.

The results revealed a significant main effect of salience-condition (F(2,406) = 18.74, p < 0.001; η²p = 0.09), a main effect of target-probability location (F(1,406) = 32.15, p < 0.001; η²p = 0.08), a main effect of BA (F(2,406) = 28.88, p < 0.001; η²p = 0.13), as well as a significant target-probability location × BA interaction (F(1,406) = 7.88, p < 0.001; η²p = 0.04). All other main effects and interactions were not significant (all p > 0.22). We investigated further the significant effects using pair-wise paired t-tests.

The comparisons between HTPL and LTPL, separately for the 3 BAs, revealed a significant effect of statistical regularities in BA17 (HTPL > LTPL, t(68) = 4.93, p < 0.001, Cohen’s d = 0.59) and BA19 (HTPL > LTPL, t(68) = 2.85, p = 0.017, Cohen’s d = 0.34), but not in BA18 (p > 0.9). In BA17 and BA19, the spatial bias vector was more distant from the target location in HTPL than in LTPL, which explains the target-probability location × BA interaction. In BA17 (top-left panel of Fig. 5), the effect of target-probability location was most evident, with the dark-colored vectors (HTPL) that are further away from the 45° dotted-line than the bright-colored vectors (LTPL). For BA19, this difference can be seen in the top-right panel of Fig. 5, again with the HTPL vectors being further away from the dotted-diagonal compared to the LTPL vectors. Thus, in BA17 and B19, presenting the target at HTPL results in a loss of the spatial bias towards the target location (i.e. the 45° dotted-line), compared with targets presented in the LTPL condition (see also dark-colored bars vs. bright-colored bars in Fig. 5, bottom panels).

**Figure 5.**
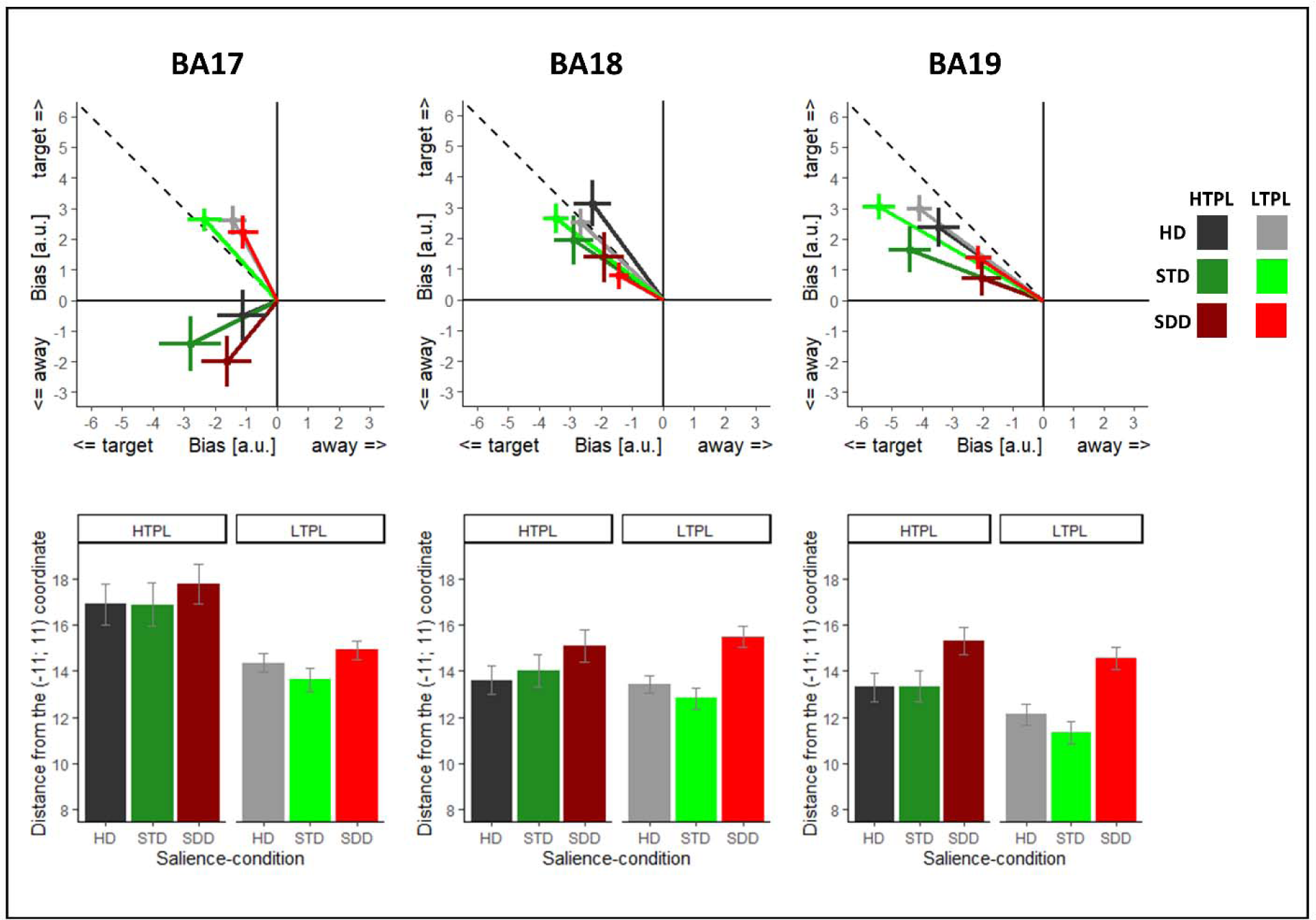
Spatial bias vectors as a function of salience-condition and target-probability location. Top panels show the 2D spatial bias vectors in BA17, BA18 and BA19 (left to right). The 45° dotted line represents the direction of the target-quadrant, in the arbitrary frame of reference of the bias vectors. The vectors represent the overall spatial bias in the occipital cortex, computed using the combined activity of the 4 ROI that represent the visual location of the 4 search items (see also Beffara et al., 2022). Thus, the distance-measure (bottom panels) reflects how far the spatial pattern of occipital activity is from an “ideal” priority map in which only the target representation would be represented. Bottom panels show mean distance (± SEM) between a point in the target quadrant (−11; 11) and the spatial bias vectors for the 2 target-probability locations and the 3 salience-conditions. Black color is used for the HD condition, green for STD and red for SDD. Darker color are vectors in HTPL condition and brighter color are vectors in LTPL condition.

Pair-wise comparisons between the 3 salience-conditions revealed that salient distractors decreased the bias towards the target location compared to both homogeneous display (i.e. higher distance values for SDD than for HD, t(137) = 8.65, p < 0.001, Cohen’s d = 0.74) and salient target display (distance values in SDD > STD (t(137) = 7.64, p < 0.001, Cohen’s d = 0.65), while salient targets did not significantly strengthen the bias towards the target location (STD > HD, p = 0.63). These results indicate that salience decreases the spatial bias towards the target location when associated with the distractor, see also results in the section above (ANOVA considering activity only in ROI_IN and ROI_DIAG).

Finally, pair-wise comparisons between the 3 BA-areas, irrespective of condition, showed that the spatial biases towards the target-quadrant increased progressively from early to higher-level visual areas (i.e. lower distance values in BA19 < BA18, t(137) = −3.04, p = 0.009, Cohen’s d = −0.26; BA18 < BA17, t(137) = 5.36, p < 0.001, Cohen’s d = −0.46, Cohen’s d = 0.46, BA19 < BA17, t(137) = −7.52, p < 0.001, Cohen’s d = −0.64), consistent with results in the section above.

In sum, we found evidence that statistical regularities modulate the spatial bias in area BA17 and BA19, with a loss of the spatial specificity towards the target location in the HTPL condition. In addition, we found that salient distractors reduced the attention bias towards the target location (SDD vs. HD, see also Beffara et al., 2022; Sprague et al., 2018). Related findings could be observed using a different set of ROIs that considered probabilistic maps of retinotopic visual areas (see Fig. S2). These results provide evidence regarding the modulation of the spatially-specific occipital responses by statistical regularities and salience. By showing vectors’ deviations away from the 45° diagonal (i.e. the target location), the bias vector analyses highlight the contribution of all the 4 regions that represent the visual space.

### 3.3. Whole-brain analyses

Beside the modulation of the activity in the occipital cortex, previous studies pointed to the parietal cortex as a key region involved in the processing of salience and search configuration probabilities (Won et al., 2020; Zhang et al., 2021 and see Geng, 2014; Geng & Duarte, 2021 for reviews). Accordingly, we performed a whole-brain group analysis using a 2×3 ANOVA (target probability location × salience-condition). The direct comparison of LTPL > HTPL revealed a significant activation in the bilateral PPC (left and right Superior Parietal Gyrus, SPG), left dorsal premotor cortex (left Superior Frontal Sulcus, SFS), right Supramarginal Gyrus as well as the anterior part of the left occipital cortex, mostly located in BA19 (see Fig. 6C). The reverse HTPL > LTPL contrast did not reveal any significant effect at the whole-brain level. When comparing SDD > STD, we found significant clusters of activation in areas of the dorsal fronto-parietal network (bilateral SFS and bilateral SPG) and in the anterior part of the left occipital cortex (see Fig. 6C and Table 1), mostly located in BA19. The reverse STD > SDD contrast revealed a significant cluster in the left Superior Frontal Sulcus (MNI x, y, z = −18 38 52, Z-value = 3.78, p = 0.042).

**Figure 6.**
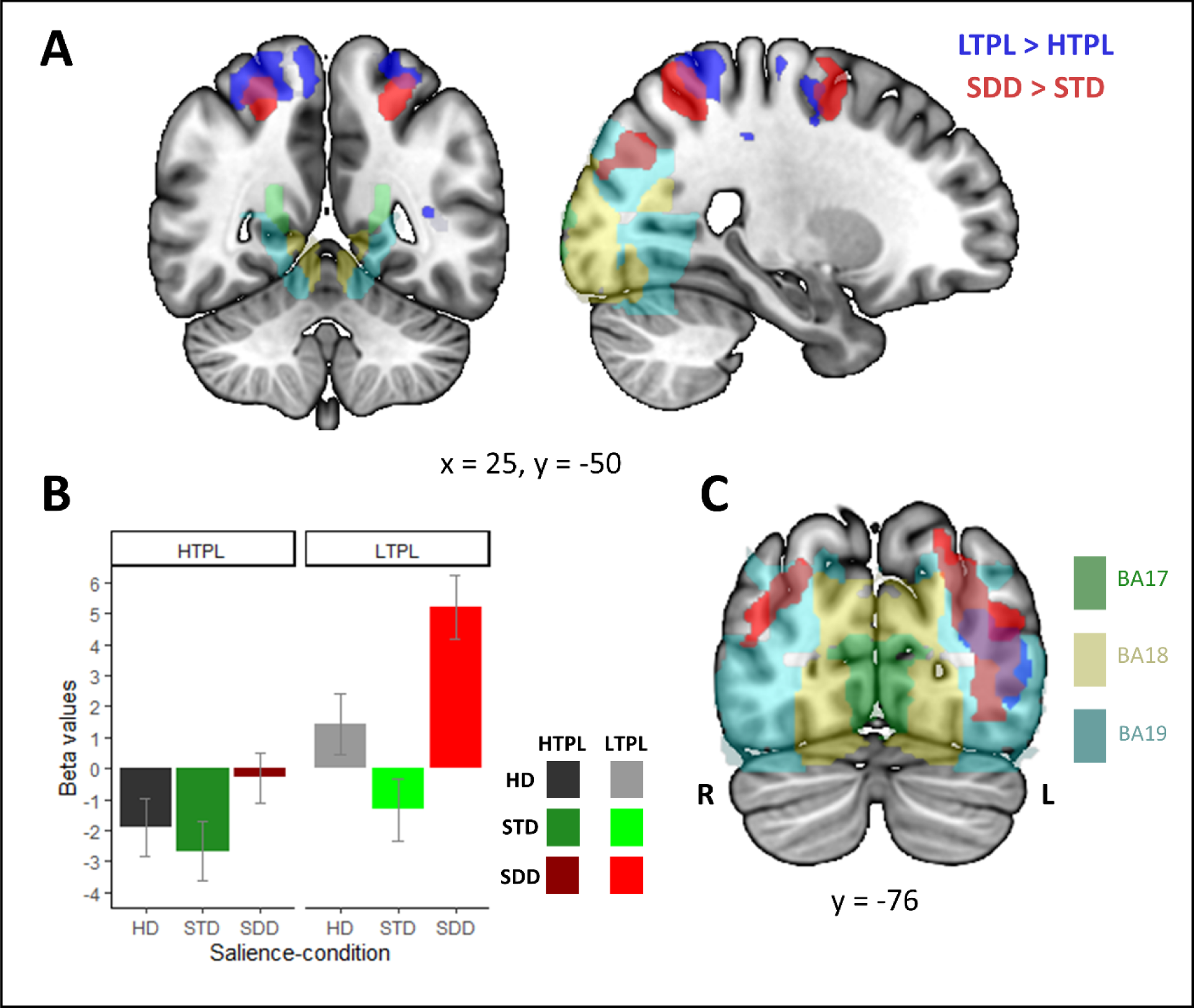
Whole-brain activations associated with salient distractors and target at low-probability locations. **A** – Coronal and sagittal sections showing the overlap between salient distractors (SDD > STD) and low target-probability location (LTPL > HTPL) in the posterior parietal cortex. **B** – Mean (± SEM) parameters estimates (beta values) for the voxels in the posterior parietal cortex showing both an effect of salience and of target-probability location (cf. region rendered in purple-color, in panel A). **C** - Occipital overlap between salience (SDD > STD) and target-probability location (LTPL > HTPL), primarily in BA19. Activation clusters are rendered a p-FWE-corr. = 0.05 at the cluster level, with cluster sizes estimated at p-unc. = 0.005.

**Table 1.** Whole-brain activations for SDD > STD and LTPL > HTPL contrasts.

| Comparison | Region | Cluster size (voxels) | Cluster-level p-value (FWE-corrected) | Z-score | MNI-coordinates |  |  |
| --- | --- | --- | --- | --- | --- | --- | --- |
| SDD > STD |  |  |  |  | x | Y | Z |
|  | L Superior Frontal Sulcus | 833 | < 0.001 | 5.37 | -22 | -2 | 48 |
|  | L Superior Frontal Sulcus | 656 | 0.001 | 4.70 | -8 | 12 | 52 |
|  | L Lateral Occipital Gyri | 2254 | < 0.001 | 4.37 | -38 | -78 | 10 |
|  | L Superior Parietal Gyrus |  |  |  | -26 | -54 | 56 |
|  | R Superior Parietal Gyrus | 426 | 0.015 | 4.10 | 24 | -52 | 54 |
|  | R Superior Frontal Sulcus | 412 | 0.018 | 3.92 | 20 | 0 | 52 |
|  | R Lateral Occipital Gyri | 531 | 0.004 | 3.70 | 38 | -66 | 22 |
| LTPL > HTPL |  |  |  |  |  |  |  |
|  | L Lateral Occipital Gyri | 1489 | < 0.001 | 4.64 | -42 | -58 | 4 |
|  | R Superior Parietal Gyrus | 1158 | < 0.001 | 4.58 | 18 | -56 | 64 |
|  | L Supramarginal Gyrus | 3466 | < 0.001 | 3.86 | -36 | -40 | 56 |
|  | L Inferior Frontal Gyrus |  |  |  | -26 | -8 | 58 |
|  | R Inferior Frontal Gyrus |  |  |  | 36 | -12 | 56 |

Thus, at the whole-brain level, we found that the processing of salience and target-probability location rely on the activation of the posterior parietal cortex (see overlap in Fig. 6, rendered in purple-color). The signal plot in Fig. 6 shows that, in the PPC, activity increased in the presence of a distractor singleton compared to displays including a salient target (SDD vs. STD, red vs. green bars). In this region, targets appearing at low-probability locations elicited greater responses than targets appearing at the high-probability location (LTPL vs. HTPL, bars 4-6 vs. bars 1-3). These results suggest that increasing the competition between spatial locations, by including distractor singletons and by presenting targets at low probability locations, increases the requirement of engaging the dorsal attention control network.

### 3.4. Psychophysiological interactions

Finally, we explored possible links between the effects of the salience-condition and the target-probability location in the PPC (cf. whole-brain analysis, and Fig. 6A), with the modulatory influences that we observed in the occipital visual cortex (see Fig. 4-5). For this, we carried out a psycho-physiological interaction analysis using the voxels in PPC that showed an overlap for the SDD > STD and the LTPL > HTPL contrasts (see Fig. 6A purple-color, and Methods Section).

Concerning the effect of salience-condition, the contrast testing for increased interregional coupling in SDD > STD at the whole-brain level did not reveal any significant effect. A more focused approach that considered specifically the coupling between PPC and the anatomically-defined occipital ROIs (BA17, BA18, BA19) did not reveal any significant effect either. The LTPL > HTPL comparison at the whole-brain level showed a significant cluster in the left middle occipital gyrus, extending ventrally to the inferior occipital gyrus (MNI x, y, z = −32, −84, 16; Z-value = 5.14, p-FWE-corrected = 0.005, n. voxels = 448). Considering more specifically the anatomically-defined occipital ROIs revealed significant effects in BA18 (MNI x, y, z = −32 −84 14, p-FWE-corrected at the voxel level = 0.012, Z-value = 4.50) and BA19 (MNI x, y, z = −32 −84 16, p-FWE-corrected at the voxel level = 0.001, Z-value = 5.14), but not in BA17 (see Fig. 7).

**Figure 7.**
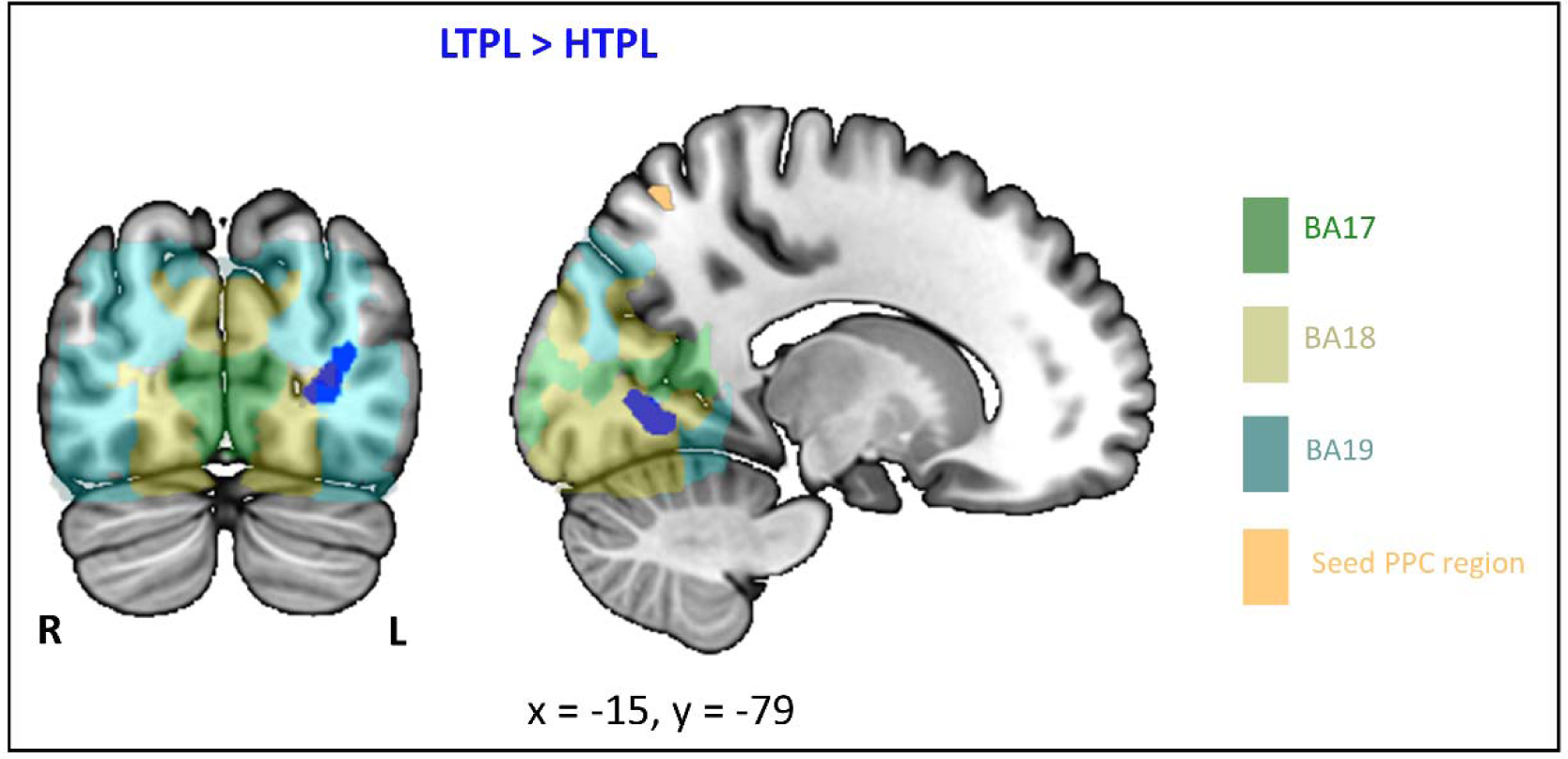
Results of the functional connectivity analyses with seed region in the PPC. The PPI analysis revealed increased connectivity between the PPC-seed region (rendered in yellow/sand color) and areas in the right occipital cortex for target presented at the low vs. high probability location (LTLP > HTPL, rendered in blue). A more focused approach that targeted anatomically-defined ROIs in the occipital cortex (small volume correction) revealed significant coupling between the PPC and voxels in BA18 and BA19, but not in BA17. The PPI effect is rendered at a p-FWE-corrected = 0.05 at the cluster level, with cluster sizes estimated at p-uncorrected = 0.005.

These additional exploratory analyses show that the target-probability location modulates the connectivity between the PPC and high-order occipital visual areas, while there was no evidence of changes of connectivity between the PPC and lower levels of the visual hierarchy (BA17). This suggests that the reduction of the spatial bias associated with the HTPL condition that we found in BA17 (cf. section 3.2.2, above) may arise via bottom-up mechanisms, without any contribution of the dorsal attention control network (here the PPC, see Discussion section).

## 4. DISCUSSION

Most previous work on signals relevant for visuo-spatial attention control has focused on the study of priority signals tested in isolation from each other (e.g. manipulation of the probability of valid/invalid cues in the classical Posner’s paradigm, Posner, 1980, see also Doricchi et al., 2010; or manipulation of target/distractor salience in Sprague et al., 2018). By contrast, in the current study we investigated how the presence of multiple attention biasing signals jointly shape visuo-spatial representations in the occipital cortex. Our results showed that both statistical regularities and sensory salience modulate spatial representations throughout the visual hierarchy, that here were operationalized using anatomical sub-divisions (BA17, BA18, BA19; cf. also Beffara et al., 2022). The whole-brain analyses revealed overlapping effects of statistical regularities (LTPL > HTPL) and salience (SDD > STD) in the posterior parietal cortex (PPC), that subsequent exploratory analyses of inter-regional connectivity further linked with the changes observed in the occipital cortex. Overall, this pattern of results indicates that statistical regularities and salience engage overlapping brain regions, but also that the underlying processing mechanisms result in different signatures at the occipital level.

### 4.1. Effects of statistical regularities and visual salience on behavioral performance

The analyses of the behavioral data showed that both statistical regularities and visual salience affected performance, with targets presented at the high-probability locations (HTPL) associated with higher accuracy and faster RTs than targets presented at low-probability locations (LTPL); and salient targets and salient distractor leading to better and poorer behavioral performance, respectively. It should be noted that in our design the salient distractors were always presented at the diagonally-opposite location compared to the target (SDD condition). In principle, the participants could have used this contingency to rapidly detect the salient distractor, and shift attention to the diagonally-opposite location to discriminate the target there. This would lead to faster and more accurate discrimination of targets in SDD compared to the “homogeneous display condition” (HD). Our behavioral results highlighted exactly the opposite pattern (better performance for HD compared with SDD), indicating that the participants did not make use of the salient distractors to guide attention towards the targets.

The results showing joint effects of salience and statistical regularities are consistent with a previous study (Ferrante et al., 2018) that also showed a reduction of search performance both by salient distractors and upon the presentation of targets at LTPL. Nonetheless, these results appear at odds with other studies that instead reported no effect of salient distractors, indicating that these can be fully suppressed (e.g. see Beffara et al., 2022; Geng, 2014; Geng & Duarte, 2021; Melloni et al., 2012; Sprague et al., 2018; Won et al., 2020). This inconsistency may be related to differences in how voluntary/strategic attention was deployed during search. In Beffara et al. (2022), the task included 100%-predictive spatial cues that were presented before the display array, so that the participants knew in advance where exactly the target would appear. Thus, the participants could strategically suppress distractors processing at the uncued locations (see also Rashal et al., 2022). In Melloni et al. (2012), there were no explicit spatial cues, but in a subset of experimental runs the participants knew whether or not the search display would contain a salient item (blocked presentation of the HD, STD or SDD condition, throughout the whole run). Indeed, the results showed better search performance in blocked-runs compared with the mixed-runs, indicative of more efficient distractor suppression when the participants could predict that the search array would include a salient distractor (Melloni et al., 2012 and see Chelazzi et al., 2019 for a review).

Unlike here, Ferrante et al. (2018) reported also a significant interaction between target-probability location and salience-condition: the increase of the RTs due to the presence of a salient distractor was larger when the target was at LTPL compared with HTPL. A possible explanation for this is that Ferrante et al.’s (2018) study included only 2 salience-conditions (HD and SDD), while here there were 3 different salience-conditions (HD, STD, SDD). With only 2 salience-conditions, the participants may learn that when the display contains a salient item, this will always be a task-irrelevant distractor. We suggest that this learning process interacts with the manipulation of target-location probability (HTPL/LTPL), yielding the observed interaction at the behavioral level. By contrast, with the inclusion of the salient target condition (3 salience-conditions, here), the presence of a salient stimulus does not provide any information about the relevance/irrelevance of the salient stimulus (target or distractor). In this situation, there could be no additional salience-related learning process and no further interaction with the main manipulation of target-location probability.

In sum, our behavioral findings demonstrate that both statistical regularities and sensory salience affect the allocation of spatial processing priorities. This indicates that in the absence of any strong endogenous drive (cf. Beffara et al., 2022; Sprague et al., 2018 who used 100% predictive cues), visual salience is not fully suppressed and does contribute to the guidance of spatial attention.

### 4.2. Occipital spatial representations are modulated according to statistical regularities and salience

The biased competition model of (visual) attention control (Desimone, 1998; Desimone & Duncan, 1995) posits that multiple visual stimuli enter competition for attentional processing resources, based on their spatial location and their behavioral relevance (sensory salience, task relevance, etc). Here we show that the activity in occipital regions that represent the stimulus location/quadrant is modulated according to the different types of attention signals. Previous neuroimaging studies have shown that activity in the occipital cortex increased when a target - compared to a distractor - is presented at the represented location (Beffara et al., 2022; Melloni et al., 2012; Sprague et al., 2018). Here, we replicated this effect of task-relevance showing that the target representation (ROI_IN) had greater activity than the distractor representation (ROI_DIAG). The effect of target selection increased progressively from BA17 to BA19 (ROI-type × BA interaction) and confirmed that the target selection bias occurs along the entire visual hierarchy (see also Beffara et al., 2022; Melloni et al., 2012; Sprague et al., 2018 for similar results).

Our main aim here was to investigate how statistical regularities and visual salience affect spatially-specific activity in the visual cortex, along with the target selection bias. On the basis of our previous work (Beffara et al., 2022), we tested how the target selection bias changed according to spatial regularities and the salience-conditions. First, we examined quadrant-specific occipital responses in the regions representing the target location (ROI_IN) and the diagonal quadrant (ROI_DIAG, where the salient distractor was presented in the SDD condition). This revealed significant interactions between ROI-type (IN/DIAG) and both salience-condition and target-probability location. Subsequent tests revealed that salience impacted most when associated with a distractor (SDD > STD, in ROI_DIAG), while statistical regularities primarily reduced activity associated with the target in the high-probability condition (LTPL > HTPL, in ROI_IN). These effects did not significantly interact with BA areas, but additional tests indicated reliable effects of statistical regularity in BA17 only.

These results demonstrate that salience and target-probability location jointly affect activity in the visual cortex, and that they do so in a spatially-specific manner. In the priority map framework, this indicates a role of the occipital cortex in attention control. The observation that current processing priorities are best accounted for by the combination of the activity associated with both target and distractors (including the effect of salience primarily expressed in the ROI_DIAG) emphasize the relevance of priority coding across distributed spatial representations (cf. also vector bias analysis below). The relative priority-weights of different locations within such representations could guide feed-forward processing, thus acting as a bottom-up source of attention control and promote processing (by higher-order regions) of signals presented at the high-priority location. However, it is important to point out that higher-order regions may themselves contribute to the setting of the priority gains, via re-entrant connectivity. The latter may also explain the finding of salience effects at the different levels of the visual hierarchy (see also Beffara et al., 2022; Melloni et al., 2012), including BA17 where salience may be represented following initial computations in higher visual areas that send feedback to lower levels of the hierarchy (Veale et al., 2017). Hence, here a combination of both feed-forward and top-down modulatory feedback is likely to determine activity in the occipital spatial representations, which therefore would participate both as a “source” and as a “site” of attention control (see also below, concerning local computations related to expectations vs. inter-regional coupling between occipital and parietal cortex).

The modulatory effects of salience and statistical regularities on the activity of the occipital cortex was also evident in the spatial bias vector analysis, which considers the contribution of all the 4 regions that represent four quadrants of the visual field. This approach was specifically conceived to assess spatial attentional priorities in terms of relative weights across occipital spatially-specific representations in the priority map framework (see also Beffara et al., 2022). The vector bias results confirmed the effect of target selection that, consistent with the ROI_IN/ROI_DIAG analysis, was found to increase progressively from BA17 to BA19, as well as the impact of salience that - again - was most prominent in the SDD condition when the display included a salient distractor (rather than a salient target). Albeit consistent with the results of the ROI_IN/ROI_DIAG analysis, the null effect for salient targets (i.e. STD vs HD comparison) was somewhat unexpected. However, it should be noted that studies in which a similar methodology was employed (Beffara et al., 2022; Melloni et al., 2012) also reported no evidence for increased bias towards the target representation for STD vs. HD in BA17 (Beffara et al., 2022) and across the visual hierarchy (Melloni et al., 2012). However, in these two studies, changes of activity/connectivity involving regions of the dorsal frontoparietal attention network were consistently observed in STD vs. HD. This further specifies that the setting of visual attentional priorities is a process that dynamically recruits several brain regions outside the occipital cortex (see Vossel et al., 2014 for a review). Indeed, several previous studies showed that activity in regions of the frontoparietal network is modulated by the presentation of salient items (see Buschman & Miller, 2007; Ibos et al., 2013 for different temporal patterns of parietal-frontal activity during target vs. salient distractor presence, Won et al., 2020; Zhang et al., 2021 for dorsal parietal activity modulation following salient distractor presentation, and Melloni et al., 2012 for frontal activity modulation by salient target presence) and that it is involved in processes facilitating target processing, including distractor suppression/target enhancement (e.g. Beffara et al., 2022; and see Chelazzi et al., 2019; Geng, 2014 for reviews).

The bias vectors analysis confirmed that also statistical regularities affect spatially-specific responses in the visual cortex. The results showed a significant interaction between statistical regularities and BA areas that subsequent tests linked to significant effects only in BA17 and BA19. In both regions, these analyses showed a loss of the attentional bias for targets presented at the high-compared to the low target-probability location (Fig. 5). The vector finding in BA17 is consistent with the results of the ROI_IN/ ROI_DIAG analysis (see above). Moreover, it also demonstrates that, not only HTPL resulted in a reduction of activity in ROI_IN, but it also modified the relationship between activity in ROI_CONTRA and ROI_IPSI. In particular, the bias-vectors in the three HTPL conditions substantially departed from the 45° diagonal (see Fig. 5, top-left panel) implying that activity in ROI_CONTRA was larger than in ROI_IPSI. An analogous effect was observed also in BA19. Overall, the bias-vectors in BA19 highlight that the attentional bias there is not strictly limited to the target quadrant representation, but comprises a more global “hemifield bias” (i.e. all 6 vectors depart from the 45° diagonal, see also Beffara et al., 2022 for analogous findings). Here we show that this shift towards a hemifield bias (as opposed to the target-quadrant bias) was stronger in the HTPL compared with the LTPL condition. As in BA17, these effects arise from a change of the relative activation of the regions representing the IPSI and CONTRA quadrants (please note that IPSI refers to “same side/hemifield” of the target quadrant). Notably, this additional contribution of ROI_CONTRA/IPSI permits detecting the effect of statistical learning in BA19, which was not significant when considering only activity in ROI_IN/DIAG (see above). The latter observation highlights the relevance of considering the distributed activity across regions representing the 4 visual quadrants to best characterise the coding of spatial processing priorities in the visual cortex (cf. also the “priority map” framework).

A note of caution should be raised here about the specific functional areas expressing the effect of statistical regularities along the visual hierarchy. Consistent with our previous work (Beffara et al., 2022), we used anatomical BAs to subdivide the quadrant-specific responses into lower-to-higher visual areas. However, these do not strictly correspond to the retinotopic areas V1/V2/V3/etc., as it can be seen in Fig. S2, panel B. An additional vector bias analysis splitting quadrant-specific responses according to a probabilistic atlas of the retinotopic areas (Wang et al., 2015) revealed significant effects of statistical regularities in V2 and V3 (see Fig. S2, panel C). The same analysis highlighted significant effects of salience in higher-order occipital regions (i.e. a ROI that comprised V3a/b, hV4, VO1/2, and LO1/2). Thus, these additional analyses confirmed the overall pattern, with an effect of statistical regularities in lower-level regions and most pronounced effects of salience at later stages, but also highlighted that precise mapping of these effects onto retinotopic areas would require identifying these areas at the participant-specific level (cf. high number of uncategorized quadrant-specific voxels in Fig. S2, panel B). The lack of these data is a limitation of the current study, which nonetheless permitted us to highlight differential effects of statistical regularities and salience along different levels of the visual hierarchy (lower/BA17, mid/BA18, higher/BA19).

The finding of a reduced bias towards the target location in the high-target probability condition is consistent with the “expectation suppression” account. This posits that responses to expected stimuli are lower than responses to unexpected stimuli (Alink & Blank, 2021; Richter & de Lange, 2019; Won et al., 2020; Zhang et al., 2021). The expectation suppression account has never been tested in the context of a spatial manipulation of the target-probability location, because previous studies focused on activity related to salient distractor stimuli only (Adam & Serences, 2021; Won et al., 2020; Zhang et al., 2021). Specifically, previous studies could not disentangle two distinct mechanisms. On the one hand, any expectation-related reduction of activity in regions that represent the distractor may indicate that expected distractors are suppressed proactively via top-down processes, so as to reduce their ability to capture attention (Adam & Serences, 2021; Geng & Duarte, 2021; Won et al., 2020; Zhang et al., 2021). But, on the other hand, the very same effect can be interpreted as evidence of sensory expectation suppression mechanisms taking place locally in the occipital cortex, related to the repeated presentation of a same stimulus (i.e. repeated presentation of the salient distractor in Adam & Serences, 2021; Won et al., 2020) at a specific location (the high-probability location of the salient distractor in Zhang et al., 2021; see Alink & Blank, 2021 for a review). Although here we did not seek to specifically address mechanisms of expectation suppression (in fact, in the priority map framework, our main prediction concerned the joint enhancement of activity associated with behaviorally-relevant spatial locations by both salience and statistical regularity), our current results show suppression at the high-probability target location and extend previous findings that considered salient distractors (Zhang et al., 2021; see also Richter & de Lange, 2019, for target-related findings but in the non-spatial domain). The latter suggests that the suppression mechanism observed in previous studies (Adam & Serences, 2021; Won et al., 2020; Zhang et al., 2021) was presumably not solely due to proactive processes, but reflected also sensory suppression mechanisms. Here, the HTPL condition led to a reduction of the spatial bias, but to an increase of the behavioral performance, suggesting a complex relation between the activity in these early sensory regions and the final behavioral outcome (cf. Alink & Blank, 2021, and see below).

### 4.3. Statistical regularities and visual salience beyond the occipital visual cortex: the contribution of the posterior parietal cortex

Although our main hypotheses concerned the modulation of quadrant-specific activity in visual cortex, we also sought to explore the possible role of other regions using whole-brain analyses and inter-regional connectivity. The main result of the whole-brain analyses was that salient distractors (SDD > STD) and targets presented at low probability locations (LTPL > HTPL) activated the posterior parietal cortex, part of the dorsal fronto-parietal attention control network (cf. Fig. 6). In a previous study, Zhang et al. (2021) found an increase of activity in the SPL when distractors were presented at an unexpected location, compared to an expected location. Similarly, Won et al. (2020) found an increased IPS activity when a salient distractor was present in the display compared to when there was no salient distractor. Our current results confirm and extend Zhang et al.’s (2021) and Won et al.’s (2020) work here showing that changes of processing properties related to the manipulation of the target - as opposed to distractors - probability location also engage the posterior parietal cortex. Further, the finding of co-occurring effects of statistical regularities and salience in the posterior parietal cortex (albeit our analyses do not imply processing by the same neural circuits) suggests that the dorsal attention network is recruited when the display configuration entails high levels of spatial competition between the current goal and other attention biasing signals (i.e., target judgment at one location, with a salient distractor and/or high-target probability at a different location). These interactions may be related to the presence of priority maps in the dorsal parietal cortex (Ptak, 2012), which would enable parallel processing of multiple, co-occurring signals. Alternatively, the activation of dorsal parietal cortex may relate to additional shifts of spatial attention that, in the context of a serial functioning of spatial control, would be required to re-orient attention from the high-target probability location to the actual target position in the LTPL condition, or from the attention-grabbing salient distractor to the non-salient, but task-relevant target in the SDD condition. While the current dataset does not enable us to exclude the latter interpretation, parallel search strategies are privileged during easy attentional tasks with few visual items (Gaspelin et al., 2023), as it was the case here, and hence we favour the former interpretation comprising competitive mechanisms between stimuli presented simultaneously.

With a set of additional, exploratory analyses of functional connectivity, we sought to link the co-occurring effects of statistical regularities and salience in the parietal cortex (cf. overlap in Fig. 6A) with our main findings in the occipital cortex. Using PPI, we assessed the functional coupling between voxels in PPC showing responses to both statistical regularities and salience, and the occipital cortex. The results showed that the coupling between PPC and BA19, as well as PPC-BA18, increased significantly in the LTPL condition (LTPL > HTPL), while this was not the case in BA17. Despite being largely preliminary, these additional observations provide us with further evidence that any suppression mechanisms taking place in the early visual cortex (V1/BA17) do not rely on interactions with the dorsal attentional network, but may rather arise via local processes (Alink & Blank, 2021). However, the significant changes of connectivity between the PPC and BA18/BA19 nuances the top-down/bottom-up dichotomy, indicating that, at least for the statistical regularity signals, the processing of attentional priorities depends on a mixture of both local computations in BA17 and occipito-parietal interactions.

Overall, the relationship between the effect of statistical regularities in the occipital cortex, its impact of the occipito-parietal connectivity, and the final behavioral outcome appears complex and multifaceted. We put forward that this arises because multiple processes are at play. Specifically, we propose that statistical regularities lead to two distinctive effects: The first one concerns the reduction of the spatial bias in BA17 when the target appears in the HTPL vs. in the LTPL. We interpret this effect in the context of predictive coding (Kok, Jehee, et al., 2012; Kok, Rahnev, et al., 2012) – here applied to the domain of spatial attention – with two apparently opposed, but synergetic effects of attentional priorities in early visual areas. On the one hand, target selection would increase the attentional gain at the target location (i.e. activity in ROI_IN > ROI_DIAG, as predicted by the biased competition model, see Desimone & Duncan, 1995). On the other hand, the positive match between expectations about the target-location and the actual target-location in HTPL would decrease activity in the same quadrant-specific representation (i.e. decreased prediction error, see Kok, Jehee, et al., 2012; Kok, Rahnev, et al., 2012). The combination of these two effects would lead to an increased precision of the target representation (“sharpening”, see Alink & Blank, 2021) when this is expected, i.e. in the HTPL condition, with the attentional gain tuned towards a better/sharpened representation of the target in the occipital cortex. This effect would primarily rely on local occipital mechanisms, thus minimizing the engagement of additional areas (i.e. top-down control for parietal cortex, cf. PPI results). This mechanism would promote target selection without taxing cognitive demands (Geng & Duarte, 2021), but would still be sufficient to affect behavior (see also Won et al., 2020). The second effect concerns the functional coupling between the PPC and the occipital regions (BA18 and BA19). The PPI results indicate that coupling increases when the target appears at a low-probability location, i.e. when attention has to be directed towards a “less obvious target location”. This may involve both feedforward and feedback signaling to “counteract” any enhanced (feedforward) processing of distractors presented at HTPL, as well as (feedback/top-down) signaling to orient attention towards the LTPL target location and boosting processing there. The two mechanisms (early local effects for HTPL targets, and PPC-mediated selection of LTPL targets) would support attentional allocation to the task-relevant target stimulus both in HTPL and LTPL conditions, but may have a different impact on the final response speed.

The analyses of inter-regional connectivity also tested whether the PPC coupling changed as a function of salience. Specifically, we tested any connectivity increase when the display contained salient distractors (SDD > STD), which in the whole-brain intra-regional analysis were found to activate the PPC. This did not reveal any significant effect in the occipital areas BA17, BA18 and BA19, where activity was modulated according to distractor salience (cf. Fig. 4-5). The lack of any salience-related inter-regional effects between PPC and the occipital cortex should not be over interpreted, but it may be consistent with the proposal that salience was processed locally. By contrast, the PPC may participate in later-stage computations that link occipital visual processing with the final behavioral output (see also above). Future studies may attempt to formally link the activity of visual regions, PPC and behavior using more complex models of effective connectivity (e.g. dynamic causal modeling, Beffara et al., 2022; Friston et al., 2003), testing for differential effects of interference on occipital vs. parietal processing (e.g. using transcranial magnetic stimulation; cf. e.g. Mevorach et al., 2006) on the behavioral patterns reported here, and/or using methods with higher temporal resolution to characterize the (feedforward/feedback) origin of occipital activity (e.g. Lamme & Roelfsema, 2000). On a final note, we must add that the current sample size is comparable with previous fMRI studies investigating spatial attention as part of the priority maps framework (number of participants ranging from 8 to 30, see Adam & Serences, 2021; Beffara et al., 2022; Melloni et al., 2012; Richter & de Lange, 2019; Sprague et al., 2018; Won et al., 2020; Zhang et al., 2021), including the ones for which power analyses have been conducted (Beffara et al., 2022; Richter & de Lange, 2019; Won et al., 2020; Zhang et al., 2021). However, larger sample sizes are of course preferable, and the field of fMRI research (Mills-Finnerty, 2021) makes no exception to the replication crisis problems (Nosek et al., 2022; Shrout & Rodgers, 2018). Therefore, future studies may be particularly cautious and assess *a priori* the required sample to detect effects of interest.

## 5. CONCLUSIONS

We investigated the joint impact of target selection, salience and target-related statistical regularities on spatial representations along the visual processing hierarchy. We found that all these attentional biasing signals affected occipital activity in a spatially-specific manner. Salient distractors’ effects were observed across the whole visual hierarchy and independently of top-down control from the dorsal fronto-parietal network. Statistical regularities decreased the spatial bias towards the target representation primarily in BA17, and this was independent of parietal-occipital interactions. However, statistical regularities affected the functional connectivity between PPC and both BA18 and BA19. These results suggest that the setting of processing priorities in the occipital cortex relies on both parietal-occipital interactions, as well as on local processing within the occipital visual cortex. These findings contribute to the debate concerning how statistical regularities affect attention control, here pointing to substantial differences compared to salience processing: while salience was associated primarily with local effects within the occipital cortex, statistical regularities engaged a combination of local occipital processing and occipito-parietal interactions.

## Data availability statement

Data available upon request.

## Ethics approval statement

The study was approved by a national ethics committee in biomedical research (Comité de Protection des Personnes: Sud-Méditerranée II, authorization ID: 2019-A00713-54).

## REFERENCES

Adam, K. C. S., & Serences, J. T. (2021). History Modulates Early Sensory Processing of Salient Distractors. The Journal of Neuroscience, 41(38), 80071–8022. 10.1523/JNEUROSCI.3099-20.2021

Alink, A., & Blank, H. (2021). Can expectation suppression be explained by reduced attention to predictable stimuli? NeuroImage, 231, 117824. 10.1016/j.neuroimage.2021.117824

Alink, A., Schwiedrzik, C. M., Kohler, A., Singer, W., & Muckli, L. (2010). Stimulus Predictability Reduces Responses in Primary Visual Cortex. Journal of Neuroscience, 30(8), 29601–2966. 10.1523/JNEUROSCI.3730-10.2010

Anderson, B. A. (2016). The attention habit1: How reward learning shapes attentional selection: The attention habit. Annals of the New York Academy of Sciences, 1369(1), 241–39. 10.1111/nyas.12957

Anderson, B. A. (2021). Time to stop calling it attentional “capture” and embrace a mechanistic understanding of attentional priority. Visual Cognition, 29(9), 5371–540. 10.1080/13506285.2021.1892894

Anderson, B. A., Laurent, P. A., & Yantis, S. (2011). Value-driven attentional capture. Proceedings of the National Academy of Sciences, 108(25), 103671–10371. 10.1073/pnas.1104047108

Awh, E., Belopolsky, A. V., & Theeuwes, J. (2012). Top-down versus bottom-up attentional control1: A failed theoretical dichotomy. Trends in Cognitive Sciences, 16(8), 4371–443. 10.1016/j.tics.2012.06.010

Bates, D., Mächler, M., Bolker, B., & Walker, S. (2015). Fitting Linear Mixed-Effects Models Using lme4. Journal of Statistical Software, 67(1). 10.18637/jss.v067.i01

Beffara, B., Hadj-Bouziane, F., Hamed, S. B., Boehler, C. N., Chelazzi, L., Santandrea, E., & Macaluso, E. (2022). Dynamic causal interactions between occipital and parietal cortex explain how endogenous spatial attention and stimulus-driven salience jointly shape the distribution of processing priorities in 2D visual space. NeuroImage, 255, 119206. 10.1016/j.neuroimage.2022.119206

Bisley, J. W., & Mirpour, K. (2019). The neural instantiation of a priority map. Current Opinion in Psychology, 29, 1081–112. 10.1016/j.copsyc.2019.01.002

Bourgeois, A., Chelazzi, L., & Vuilleumier, P. (2016). How motivation and reward learning modulate selective attention. In Progress in Brain Research (Vol. 229, p. 3251–342). Elsevier. 10.1016/bs.pbr.2016.06.004

Buschman, T. J., & Miller, E. K. (2007). Top-Down Versus Bottom-Up Control of Attention in the Prefrontal and Posterior Parietal Cortices. Science, 315(5820), 18601–1862. 10.1126/science.1138071

Chelazzi, L., Della Libera, C., Sani, I., & Santandrea, E. (2011). Neural basis of visual selective attention1: Visual selective attention. Wiley Interdisciplinary Reviews: Cognitive Science, 2(4), 3921–407. 10.1002/wcs.117

Chelazzi, L., Estocinova, J., Calletti, R., Lo Gerfo, E., Sani, I., Della Libera, C., & Santandrea, E. (2014). Altering Spatial Priority Maps via Reward-Based Learning. Journal of Neuroscience, 34(25), 85941–8604. 10.1523/JNEUROSCI.0277-14.2014

Chelazzi, L., Marini, F., Pascucci, D., & Turatto, M. (2019). Getting rid of visual distractors1: The why, when, how, and where. Current Opinion in Psychology, 29, 1351–147. 10.1016/j.copsyc.2019.02.004

Chelazzi, L., Perlato, A., Santandrea, E., & Della Libera, C. (2013). Rewards teach visual selective attention. Vision Research, 85, 581–72. 10.1016/j.visres.2012.12.005

Chica, A. B., Bartolomeo, P., & Valero-Cabre, A. (2011). Dorsal and Ventral Parietal Contributions to Spatial Orienting in the Human Brain. Journal of Neuroscience, 31(22), 81431–8149. 10.1523/JNEUROSCI.5463-10.2010

Clark, C. W. (2003). The behavioral ecology of a cognitive constraint1: Limited attention. Behavioral Ecology, 14(2), 1511–156. 10.1093/beheco/14.2.151

Corbetta, M., Kincade, J. M., Ollinger, J. M., McAvoy, M. P., & Shulman, G. L. (2000). Voluntary orienting is dissociated from target detection in human posterior parietal cortex. Nature Neuroscience, 3(3), 2921–297. 10.1038/73009

Corbetta, M., & Shulman, G. L. (2002). Control of goal-directed and stimulus-driven attention in the brain. Nature Reviews Neuroscience, 3(3), 2011–215. 10.1038/nrn755

Desimone, R. (1998). Visual attention mediated by biased competition in extrastriate visual cortex. Philosophical Transactions of the Royal Society of London. Series B: Biological Sciences, 353(1373), 12451–1255. 10.1098/rstb.1998.0280

Desimone, R., & Duncan, J. (1995). Neural mechanisms of selective visual attention. Annual Review of Neuroscience, 18, 1931–222. 10.1146/annurev.ne.18.030195.001205

Doricchi, F., Macci, E., Silvetti, M., & Macaluso, E. (2010). Neural Correlates of the Spatial and Expectancy Components of Endogenous and Stimulus-Driven Orienting of Attention in the Posner Task. Cerebral Cortex, 20(7), 15741–1585. 10.1093/cercor/bhp215

Dukas, R. (2004). Causes and Consequences of Limited Attention. Brain, Behavior and Evolution, 63(4), 1971–210. 10.1159/000076781

Egeth, H. E., & Yantis, S. (1997). VISUAL ATTENTION1: Control, Representation, and Time Course. Annual Review of Psychology, 48(1), 2691–297. 10.1146/annurev.psych.48.1.269

Ferrante, O., Patacca, A., Di Caro, V., Della Libera, C., Santandrea, E., & Chelazzi, L. (2018). Altering spatial priority maps via statistical learning of target selection and distractor filtering. Cortex, 102, 671–95. 10.1016/j.cortex.2017.09.027

Friston, K. J., Buechel, C., Fink, G. R., Morris, J., Rolls, E., & Dolan, R. J. (1997). Psychophysiological and Modulatory Interactions in Neuroimaging. NeuroImage, 6(3), 2181–229. 10.1006/nimg.1997.0291

Friston, K. J., Harrison, L., & Penny, W. (2003). Dynamic causal modelling. NeuroImage, 19(4), 12731–1302. 10.1016/S1053-8119(03)00202-7

Gaspelin, N., Egeth, H. E., & Luck, S. J. (2023). A Critique of the Attentional Window Account of Capture Failures. Journal of Cognition, 6(1), 39. 10.5334/joc.270

Gaspelin, N., Leonard, C. J., & Luck, S. J. (2017). Suppression of overt attentional capture by salient-but-irrelevant color singletons. Attention, Perception, & Psychophysics, 79(1), 451–62. 10.3758/s13414-016-1209-1

Geng, J. J. (2014). Attentional Mechanisms of Distractor Suppression. Current Directions in Psychological Science, 23(2), 1471–153. 10.1177/0963721414525780

Geng, J. J., & Duarte, S. E. (2021). Unresolved issues in distractor suppression1: Proactive and reactive mechanisms, implicit learning, and naturalistic distraction. Visual Cognition, 29(9), 6081–613. 10.1080/13506285.2021.1928806

Gibson, B. S., & Kingstone, A. (2006). Visual Attention and the Semantics of Space1: Beyond Central and Peripheral Cues. Psychological Science, 17(7), 6221–627. 10.1111/j.1467-9280.2006.01754.x

Gottlieb, J. (2007). From Thought to Action1: The Parietal Cortex as a Bridge between Perception, Action, and Cognition. Neuron, 53(1), 91–16. 10.1016/j.neuron.2006.12.009

Ibos, G., Duhamel, J.-R., & Ben Hamed, S. (2013). A Functional Hierarchy within the Parietofrontal Network in Stimulus Selection and Attention Control. Journal of Neuroscience, 33(19), 83591–8369. 10.1523/JNEUROSCI.4058-12.2013

Itti, L., & Koch, C. (2001). Computational modelling of visual attention. Nature Reviews Neuroscience, 2(3), 1941–203. 10.1038/35058500

Klein, J. T., Shepherd, S. V., & Platt, M. L. (2009). Social Attention and the Brain. Current Biology, 19(20), R9581–R962. 10.1016/j.cub.2009.08.010

Kok, P., Jehee, J. F. M., & de Lange, F. P. (2012). Less Is More1: Expectation Sharpens Representations in the Primary Visual Cortex. Neuron, 75(2), 2651–270. 10.1016/j.neuron.2012.04.034

Kok, P., Rahnev, D., Jehee, J. F. M., Lau, H. C., & De Lange, F. P. (2012). Attention Reverses the Effect of Prediction in Silencing Sensory Signals. Cerebral Cortex, 22(9), 21971–2206. 10.1093/cercor/bhr310

Lamme, V. A. F., & Roelfsema, P. R. (2000). The distinct modes of vision offered by feedforward and recurrent processing. Trends in Neurosciences, 23(11), 5711–579. 10.1016/S0166-2236(00)01657-X

Luck, S. J., Gaspelin, N., Folk, C. L., Remington, R. W., & Theeuwes, J. (2021). Progress toward resolving the attentional capture debate. Visual Cognition, 29(1), 11–21. 10.1080/13506285.2020.1848949

McLaren, D. G., Ries, M. L., Xu, G., & Johnson, S. C. (2012). A generalized form of context-dependent psychophysiological interactions (gPPI)1: A comparison to standard approaches. NeuroImage, 61(4), 12771–1286. 10.1016/j.neuroimage.2012.03.068

Melloni, L., van Leeuwen, S., Alink, A., & Müller, N. G. (2012). Interaction between Bottom-up Saliency and Top-down Control1: How Saliency Maps Are Created in the Human Brain. Cerebral Cortex, 22(12), 29431–2952. 10.1093/cercor/bhr384

Mevorach, C., Humphreys, G. W., & Shalev, L. (2006). Opposite biases in salience-based selection for the left and right posterior parietal cortex. Nature Neuroscience, 9(6), 7401–742. 10.1038/nn1709

Mills-Finnerty, C. (2021). Five best practices for fMRI research1: Towards a biologically grounded understanding of mental phenomena. Journal for Reproducibility in Neuroscience, 2, 1517. 10.31885/jrn.2.2021.1517

Nosek, B. A., Hardwicke, T. E., Moshontz, H., Allard, A., Corker, K. S., Dreber, A., Fidler, F., Hilgard, J., Kline Struhl, M., Nuijten, M. B., Rohrer, J. M., Romero, F., Scheel, A. M., Scherer, L. D., Schönbrodt, F. D., & Vazire, S. (2022). Replicability, Robustness, and Reproducibility in Psychological Science. Annual Review of Psychology, 73, 7191–748. 10.1146/annurev-psych-020821-114157

Pessoa, L. (2010). Embedding reward signals into perception and cognition. Frontiers in Neuroscience, 4. 10.3389/fnins.2010.00017

Pessoa, L. (2015). Multiple influences of reward on perception and attention. Visual Cognition, 23(11–2), 2721–290. 10.1080/13506285.2014.974729

Posner, M. I. (1980). Orienting of Attention. Quarterly Journal of Experimental Psychology, 32(1), 31–25. 10.1080/00335558008248231

Ptak, R. (2012). The Frontoparietal Attention Network of the Human Brain1: Action, Saliency, and a Priority Map of the Environment. The Neuroscientist, 18(5), 5021–515. 10.1177/1073858411409051

Rashal, E., Senoussi, M., Santandrea, E., Ben-Hamed, S., Macaluso, E., Chelazzi, L., & Boehler, C. N. (2022). An EEG study of the combined effects of top-down and bottom-up attentional selection under varying task difficulty. Psychophysiology. 10.1111/psyp.14002

Richter, D., & de Lange, F. P. (2019). Statistical learning attenuates visual activity only for attended stimuli. eLife, 8, e47869. 10.7554/eLife.47869

Shrout, P. E., & Rodgers, J. L. (2018). Psychology, Science, and Knowledge Construction1: Broadening Perspectives from the Replication Crisis. Annual Review of Psychology, 69(1), 4871–510. 10.1146/annurev-psych-122216-011845

Sprague, T. C., Itthipuripat, S., Vo, V. A., & Serences, J. T. (2018). Dissociable signatures of visual salience and behavioral relevance across attentional priority maps in human cortex. Journal of Neurophysiology, 119(6), 21531–2165. 10.1152/jn.00059.2018

Thompson, K. G., & Bichot, N. P. (2005). A visual salience map in the primate frontal eye field. In Progress in Brain Research (Vol. 147, p. 2491–262). Elsevier. 10.1016/S0079-6123(04)47019-8

Todd, R. M., & Manaligod, M. G. M. (2018). Implicit guidance of attention1: The priority state space framework. Cortex, 102, 1211–138. 10.1016/j.cortex.2017.08.001

Tzourio-Mazoyer, N., Landeau, B., Papathanassiou, D., Crivello, F., Etard, O., Delcroix, N., Mazoyer, B., & Joliot, M. (2002). Automated Anatomical Labeling of Activations in SPM Using a Macroscopic Anatomical Parcellation of the MNI MRI Single-Subject Brain. NeuroImage, 15(1), 2731–289. 10.1006/nimg.2001.0978

Veale, R., Hafed, Z. M., & Yoshida, M. (2017). How is visual salience computed in the brain? Insights from behaviour, neurobiology and modelling. Philosophical Transactions of the Royal Society B: Biological Sciences, 372(1714), 20160113. 10.1098/rstb.2016.0113

Vossel, S., Geng, J. J., & Fink, G. R. (2014). Dorsal and Ventral Attention Systems1: Distinct Neural Circuits but Collaborative Roles. The Neuroscientist, 20(2), 1501–159. 10.1177/1073858413494269

Vossel, S., Weidner, R., Driver, J., Friston, K. J., & Fink, G. R. (2012). Deconstructing the Architecture of Dorsal and Ventral Attention Systems with Dynamic Causal Modeling. Journal of Neuroscience, 32(31), 106371–10648. 10.1523/JNEUROSCI.0414-12.2012

Vuilleumier, P. (2005). How brains beware1: Neural mechanisms of emotional attention. Trends in Cognitive Sciences, 9(12), 5851–594. 10.1016/j.tics.2005.10.011

Wang, L., Mruczek, R. E. B., Arcaro, M. J., & Kastner, S. (2015). Probabilistic Maps of Visual Topography in Human Cortex. Cerebral Cortex, 25(10), 39111–3931. 10.1093/cercor/bhu277

Wolfe, J. M., Butcher, S. J., Lee, C., & Hyle, M. (2003). Changing your mind1: On the contributions of top-down and bottom-up guidance in visual search for feature singletons. Journal of Experimental Psychology: Human Perception and Performance, 29(2), 4831–502. 10.1037/0096-1523.29.2.483

Won, B.-Y., Forloines, M., Zhou, Z., & Geng, J. J. (2020). Changes in visual cortical processing attenuate singleton distraction during visual search. Cortex, 132, 3091–321. 10.1016/j.cortex.2020.08.025

Worsley, K. J., Marrett, S., Neelin, P., Vandal, A. C., Friston, K. J., & Evans, A. C. (1996). A unified statistical approach for determining significant signals in images of cerebral activation. Human Brain Mapping, 4(1), 581–73. 10.1002/(SICI)1097-0193(1996)4:1<58::AID-HBM4>3.0.CO;2-O

Zhang, B., Weidner, R., Allenmark, F., Bertleff, S., Fink, G. R., Shi, Z., & Müller, H. J. (2021). Statistical Learning of Frequent Distractor Locations in Visual Search Involves Regional Signal Suppression in Early Visual Cortex. Cerebral Cortex, bhab377. 10.1093/cercor/bhab377

