## Supplementary material for "Separate and overlapping mechanisms of statistical regularities and salience processing in the occipital cortex and dorsal attention network": Fig. S

**Supplementary *Figure S1.***

***The main steps of the spatial bias computation.*** *The first step of the computation of the spatial bias vectors consisted in projecting the mean beta values of the 4 quadrant-specific ROIs (i.e. the four quadrant representations: top-left = TL; top-right = TR; bottom-right = BR; bottom-left = BL) in a 2D-plane. This was done separately for each of the 24 conditions (see details of the GLM analysis, in the main text). For each condition, the 4 mean betas values were used to define four orthogonal vectors along the four 45° diagonals. The length of each vector was proportional to the mean beta value of one ROI, because the vectors extended from the position (0, 0) to the position (betaROI, betaROI) in the respective quadrant. This is shown in the central panel of the figure: the mean of the betas of the region ROI_TL (blue ROI in the illustration on the top-left, and corresponding mean beta value in the bar plot below) was transformed in a vector with (0, 0) as the starting coordinate and (-betaROI_TL, betaROI_TL) as the end-point coordinate (see vector displayed in blue in the central 2D-space representation). This procedure was repeated for the mean beta values of the other 3 ROIs, using the corresponding end-point coordinates: (betaROI_BR, -betaROI_BR) (in cyan); (betaROI_TR, betaROI_TR) (in yellow); and (-betaROI_BL, -betaROI_BL) (in magenta). The 4 resulting vectors were then summed in order to obtain a single vector per experimental condition (4 target-quadrants x 2 target-probability locations x 3 salience-conditions), BA and participant (cf. bottom-right panel, illustrating the summed vector for one condition). The second step of the computation consisted in projecting the vectors associated with targets presented in the 4 quadrants in the top-left quadrant (i.e. as if all targets had appeared in the top-left quadrant, for illustrative purposes, Fig. S1 bottom-left, red arrow). The projected vectors (dotted lines in the left-bottom panel) were then averaged so as to obtain a single vector (black line in the left-bottom panel) that characterizes the spatial bias across the 4 areas that represents the 4 quadrants (BL, TL, BR, TR, in the top-left panel), irrespective of the actual quadrant-location of the target. The resulting spatial bias vectors for each target-probability location (HTPL, LTPL) and salience-condition (HD, STD, SDD) and BA area (BA17, 18, and 19) were then compared as described in the main text (see also Fig. 5).*

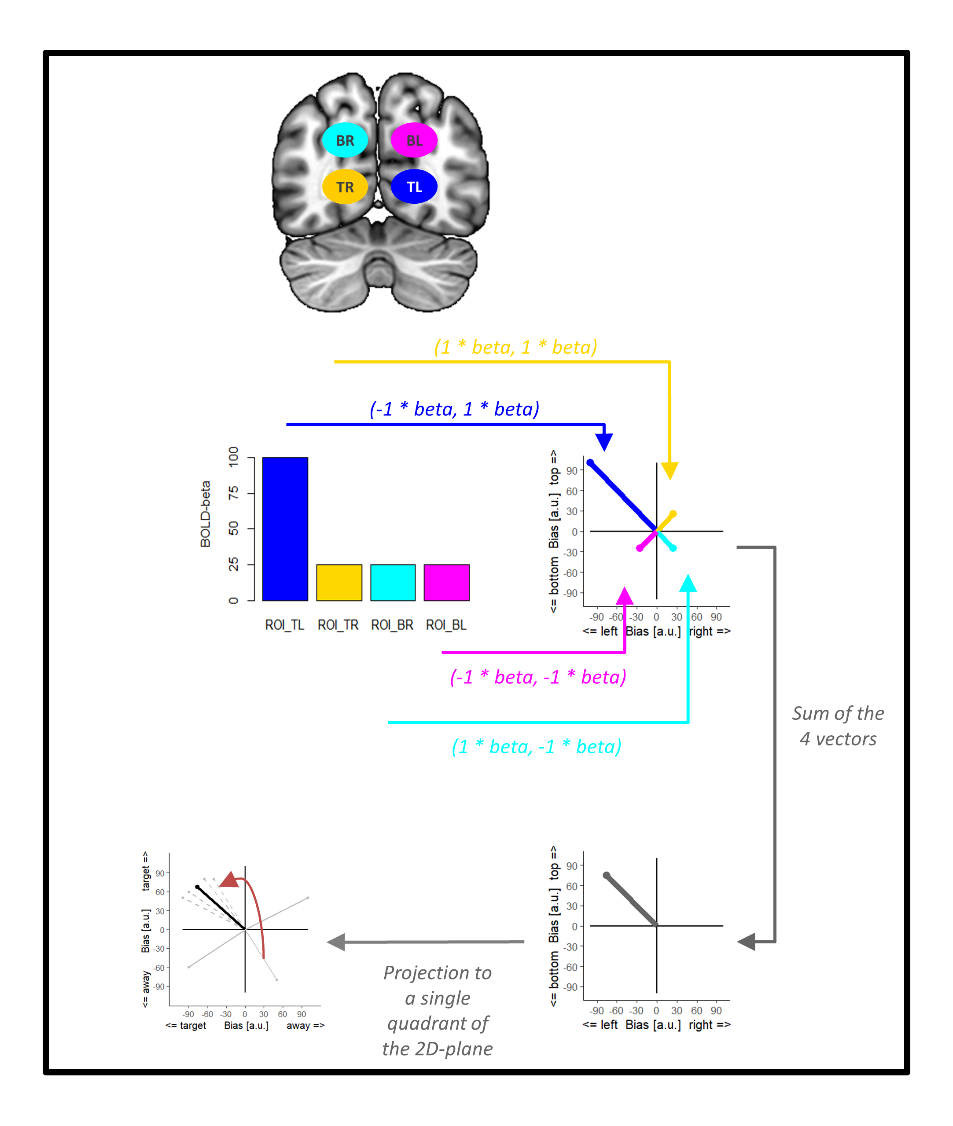

**Supplementary *Figure S2.***

***Additional analyses considering probabilistic retinotopic maps.*** ***Panel A****. Coronal sections illustrating the overlap between our functional ROI_BA17 and ROI_BA18 (which combined subject-specific quadrant mapping and anatomical masks considering Brodmann areas BA17 and BA18; Tzourio-Mazoyer et al., 2002, see also Methods section) and probabilistic maps of retinotopic areas V1 and V2 according to Wang et al. (2015). This shows a reasonable overlap between ROI_BA17 and V1: 93% of the voxels of our ROI_BA17 had a probability larger than zero to be in area V1. The match was weaker between our ROI_BA18 and retinotopic V2: 59% of the ROI_BA18 voxels had a probability larger than zero to be in area V2.* ***Panel B.*** *The bar graph shows how the voxels of our ROIs are categorized according to the retinotopic areas, now - for each voxel - considering the atlas area with the highest probability (i.e. using the "Maximum Probability Map" [MPM] of Wang et al.'s, 2015 atlas). The bar plot shows that V1 is most represented in our ROI_BA17 and V2 is most represented in our ROI_BA18, but it also indicates that each of our ROIs (and particularly so ROI_BA19) included voxels belonging to many different regions. In addition, it should be noted that the "Maximum Probability Map" was computed on the gray-matter surface and it is not continuous in the 3D volume space (see Fig. 3C in Wang et al., 2015). As a consequence of this, a large number of our ROIs’ voxels are not classified to any of the retinotopic regions (see 3 leftmost bars on the plot, and please note that the y-axis is not linear on this graph).* ***Panel C.*** *Results of a supplementary analysis of the main attention dataset for which the spatial bias vectors were now computed based on ROIs that combined our quadrant-mapping localizer data with the "Maximum Probability Map" [MPM] of Wang's retinotopic atlas. Accordingly, we defined a new set of ROIs categorizing voxels as belonging to areas V1, V2, V3 and pooled higher-order occipital areas in single ROI (i.e. "ROI_higherV", which included the other dorsal and ventral occipital areas: V3a/b, hV4, VO1/2, and LO1/2). These new analyses revealed a pattern overall consistent with our main analyses: HTPL reduced the bias towards the target position at relatively early stages (statistically significant in ROI_V2, F(2,110) = 5.6, p < 0.021; and ROI_V3, F(2,110) = 4.9, p < 0.029), while the effect of salience was most pronounced at later stages (statistically significant in ROI_ higherV; F(2,110) = 11.2, p < 0.001). As expected, the mapping of these effects to the different occipital areas was different compared to our main analysis, because: 1) the match between BAs and retinotopic areas is relatively poor; and 2) a large number of voxels responding to our quadrant-localizer were not classified using Wang et al.'s, 2015 "Maximum Probability Map", cf. panel B.*

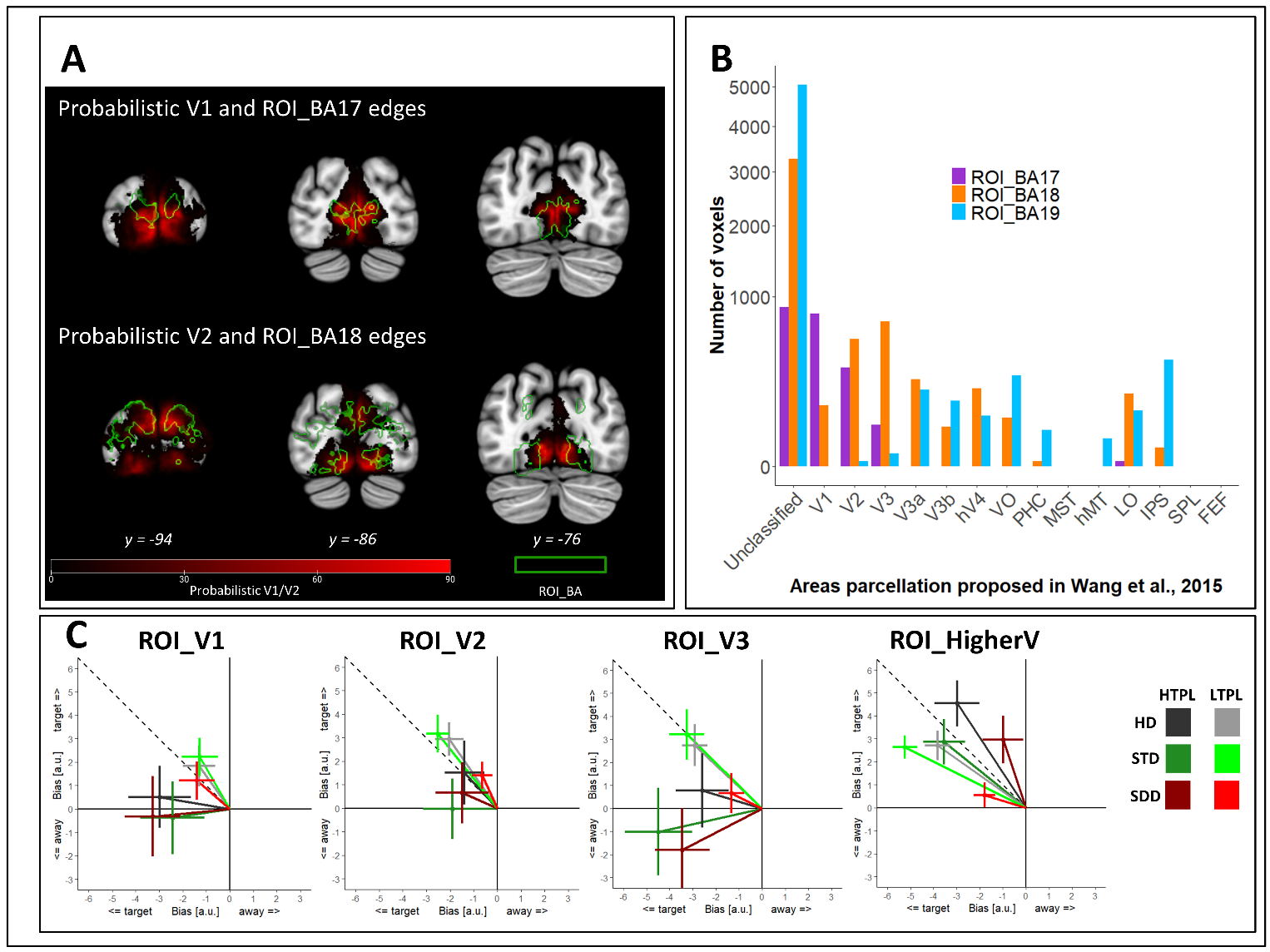

References

Tzourio-Mazoyer, N., Landeau, B., Papathanassiou, D., Crivello, F., Etard, O., Delcroix, N., Mazoyer, B., & Joliot, M. (2002). Automated Anatomical Labeling of Activations in SPM Using a Macroscopic Anatomical Parcellation of the MNI MRI Single-Subject Brain. NeuroImage, 15(1), 273‑289. https://doi.org/10.1006/nimg.2001.0978

Wang, L., Mruczek, R. E. B., Arcaro, M. J., & Kastner, S. (2015). Probabilistic Maps of Visual Topography in Human Cortex. Cerebral Cortex, 25(10), 3911‑3931. https://doi.org/10.1093/cercor/bhu277
